# Experimentally calibrated computational modeling of inflammation and vascular remodeling in breast cavity healing following breast-conserving surgery

**DOI:** 10.64898/2026.07.18.739352

**Authors:** Zachary Harbin, Carla Fisher, Rachel A. Morrison, Hector Gomez, Sherry Voytik-Harbin, Adrian Buganza Tepole

**Affiliations:** School of Mechanical Engineering, Purdue University, West Lafayette, IN, USA; Division of Breast Surgery, Indiana University School of Medicine, Indianapolis, IN, USA; Weldon School of Biomedical Engineering, Purdue University, West Lafayette, IN, USA; Department of Basic Medical Sciences, Purdue University, West Lafayette, IN, USA

**Keywords:** Mechanobiological modeling, Breast-conserving surgery, Wound healing, Vascular remodeling, Inflammatory signaling, Angiogenesis, Bayesian calibration, Multi-task Gaussian processes, Nonlinear finite elements, Growth, remodeling

## Abstract

Breast-conserving surgery (BCS; lumpectomy) is widely used to treat early-stage breast cancer, yet the complexity and patient-specific variability of postoperative cavity remodeling make healing trajectories and physical outcomes difficult to predict. Although inflammatory and vascular processes are central to these outcomes, mathematical models of tissue healing do not capture their coupled interactions or calibrate them against experimental data. Here, we extend a computational model of breast cavity healing to incorporate coupled inflammatory and vascular dynamics, including angiogenesis, oxygen transport, and inflammatory cell activity. The model is calibrated using preclinical porcine lumpectomy histology and literature data. Model parameters are inferred using a multi-task Gaussian process surrogate within a Bayesian inference framework to align predictions with experimental observations and quantify uncertainty. Thus, this work provides a mechanistically grounded framework for inflammatory and vascular remodeling during breast cavity healing and provides a foundation for patient-specific prediction of healing and physical outcomes following lumpectomy.

## INTRODUCTION

Breast cancer accounts for approximately one-third of new cancer diagnoses in women, making it one of the most prevalent and clinically significant cancers^1^. Improvements in early detection and treatment have significantly increased breast cancer survival over time, leading to a growing population of over 4 million survivors in the United States alone. This shift has placed greater emphasis on long-term outcomes, including recurrence prevention and patient quality of life^2^. Surgical intervention remains central to breast cancer treatment, as it is associated with the lowest recurrence rates^3,4^ and is most commonly performed through either breast-conserving surgery (BCS; otherwise known as lumpectomy) or mastectomy (complete removal of the breast). The choice between these surgical approaches is often complex and emotionally challenging, involving patient preferences, clinical factors, and anticipated post-surgical outcomes. In recent years, BCS has emerged as the preferred treatment for early-stage breast cancer, with approximately 60% of patients undergoing lumpectomy^5^ due to its comparable survival and recurrence outcomes, lower risk of complications, and improved preservation of breast appearance^6–10^. The primary goal of BCS is to achieve complete tumor resection while preserving healthy breast tissue and overall breast appearance. However, tumor excision creates a residual cavity that undergoes a dynamic healing response (Figure 1), often resulting in tissue contraction, scar formation, and breast deformation, including asymmetry and surface irregularities. Achieving favorable cosmetic outcomes is therefore a key objective in surgical planning, given its strong association with patient satisfaction and long-term quality of life^11,12^. Yet, predicting post-surgical healing trajectories and the resulting physical outcomes remains challenging due to the multifactorial cavity healing process.

**Figure 1:**
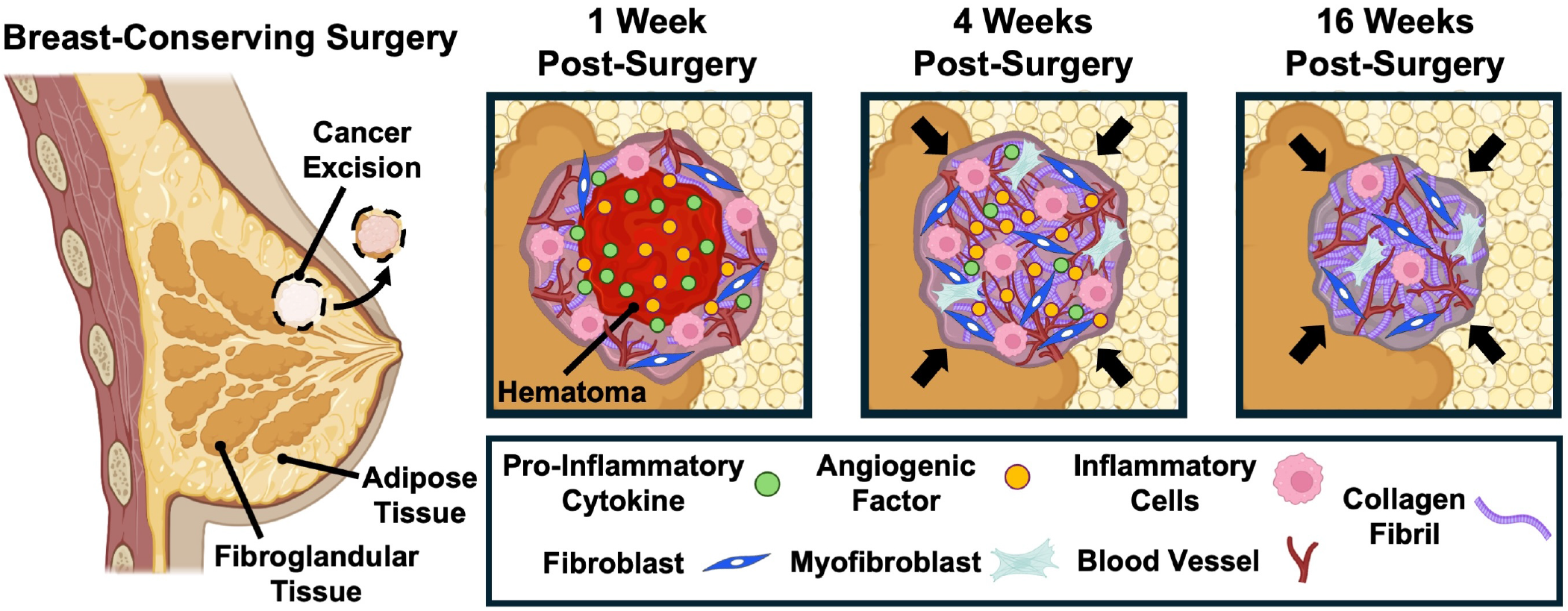
Breast-Conserving Surgery and Subsequent Cavity Healing Progression. Tumor excision creates a surgical cavity often accompanied by formation of a hematoma and/or seroma, within which a provisional fibrin matrix forms. Inflammatory cells accumulate primarily along the cavity boundary and release pro-inflammatory cytokines and angiogenic factors that establish signaling gradients promoting angiogenesis and subsequent fibroblast recruitment. The developing vasculature restores oxygen and nutrient delivery, supporting fibroblast proliferation and collagen deposition. Fibroblast differentiation into contractile myofibroblasts drives cavity contraction (indicated by black arrows) and collagen remodeling, while the vascular network gradually undergoes pruning as healing progresses. Over time, this tissue reorganizes to form dense, contracted scar tissue.

The progression of cavity healing following BCS occurs through the overlapping phases of hemostasis, inflammation, proliferation, and tissue remodeling, as illustrated in Figure 1 and informed by porcine lumpectomy study observations^13^. Immediately following tumor excision, the resulting surgical cavity fills with blood (hematoma) or serous fluid (seroma). A provisional fibrin matrix forms within the cavity, providing a temporary scaffold for tissue repair. Disruption of tissue continuity and perfusion creates hypoxic conditions that contribute to inflammatory cell recruitment and initiation of angiogenesis. This environment drives the recruitment of inflammatory cells, including macrophages, which localize primarily along the cavity boundary, while the cavity center remains dominated by fluid with minimal cellular presence. Macrophages at the cavity-tissue interface release pro-inflammatory cytokines and angiogenic factors that establish signaling gradients promoting angiogenesis. With increasing vascularization and improved oxygen delivery, fibroblast proliferation and migration are supported, while remodeling of the provisional matrix and collagen deposition lead to the formation of granulation tissue. Fibroblasts subsequently differentiate into myofibroblasts, generating contractile forces that further drive cavity contraction and alignment of the collagen matrix. As the inflammatory response subsides, the developing vascular network undergoes maturation and pruning, while the collagen matrix continues to reorganize, ultimately forming a dense, fibrotic scar. These structural changes lead to and altered breast consistency and shape, adversely impacting patient quality of life^14^.

Computational modeling offers a mechanistic framework for linking cellular, biochemical, and biomechanical processes to wound healing outcomes. Most efforts have focused on cutaneous (skin) wound healing, where models have progressively evolved to capture increasing levels of biological complexity, including cellular activity and tissue remodeling^15–25^. Yet, there is still a gap in models that capture the coupling between i.nflammatory activity, angiogenic signaling, oxygen delivery, cellular recruitment, and matrix remodeling. Moreover, many models rely on idealized dynamics rather than direct calibration to experimental observations. Similar limitations apply to the few modeling approaches that have been applied to breast cavity healing following BCS^26–28^.

Previous work by our group has led to the development and calibration of a computational mechanobiological model of breast cavity healing following BCS^29,30^. The model is implemented within a finite element framework that integrates nonlinear tissue mechanics with cellular and extracellular matrix (ECM) processes governing cavity healing. This framework captures the dynamic coupling between tissue mechanics and evolving biological fields, including pro-inflammatory cytokine signaling, fibroblast activity, and collagen deposition, linking these processes to changes in tissue structure and mechanical behavior. Model parameters were informed and calibrated using a combination of porcine lumpectomy data and published clinical observations, enabling the model to reproduce observed trends in tissue remodeling and contraction^29,30^.

Despite these advances, key inflammatory and vascular processes that regulate tissue healing are not explicitly represented in the prior framework. In the context of BCS, these processes play a central role in coordinating healing and are influenced by therapeutic interventions and patient-specific clinical and physiologic factors. For example, radiation therapy, which is commonly administered following BCS to minimize cancer recurrence^31,32^, can significantly alter inflammatory and vascular responses, leading to fibrosis and exacerbating long-term tissue deformation^33–35^. In addition, comorbidities such as diabetes, as well as conditions associated with impaired perfusion, can further disrupt vascularization and alter the progression of healing and remodeling^36,37^. Therapeutics such as regenerative breast tissue fillers that promote cellularization and vascularization to support tissue repair further emphasize the importance of accurately representing these processes^13^. Collectively, these considerations motivate extension of the computational framework to more comprehensively quantify the influence of inflammatory and vascular processes on healing variability following BCS.

In this study, we extend the computational model of post-BCS cavity healing^29,30^ to incorporate key inflammatory and vascular processes by introducing coupled biochemical species governing vascular remodeling and inflammatory signaling. Model behavior is informed by experimental histology from the porcine lumpectomy model and relevant literature data to characterize the spatiotemporal evolution of inflammatory and vascular processes during healing. To align model predictions with these observations, a machine learning surrogate model is employed within a Bayesian inference framework to calibrate model parameters, reproduce experimentally observed trends, and quantify model uncertainty. Together, this work establishes a calibrated computational framework for coupled inflammatory and vascular processes following BCS and provides a mechanistically grounded foundation for future patient-specific prediction of surgical outcomes and personalized treatment planning.

## RESULTS

### Computational Model Implementation

The computational model describes the spatiotemporal dynamics of key molecular and cellular species involved in cavity healing, including fibroblasts, pro-inflammatory cytokines, collagen, oxygen, angiogenic factors, macrophages, and capillaries. These species are coupled through a system of partial (PDEs) and ordinary (ODEs) differential equations. The regulatory relationships between the modeled species are shown in Figure 2, which depicts the coupling structure and their interdependencies. A detailed description of the model equations and parameterization is provided in the STAR Methods.

**Figure 2:**
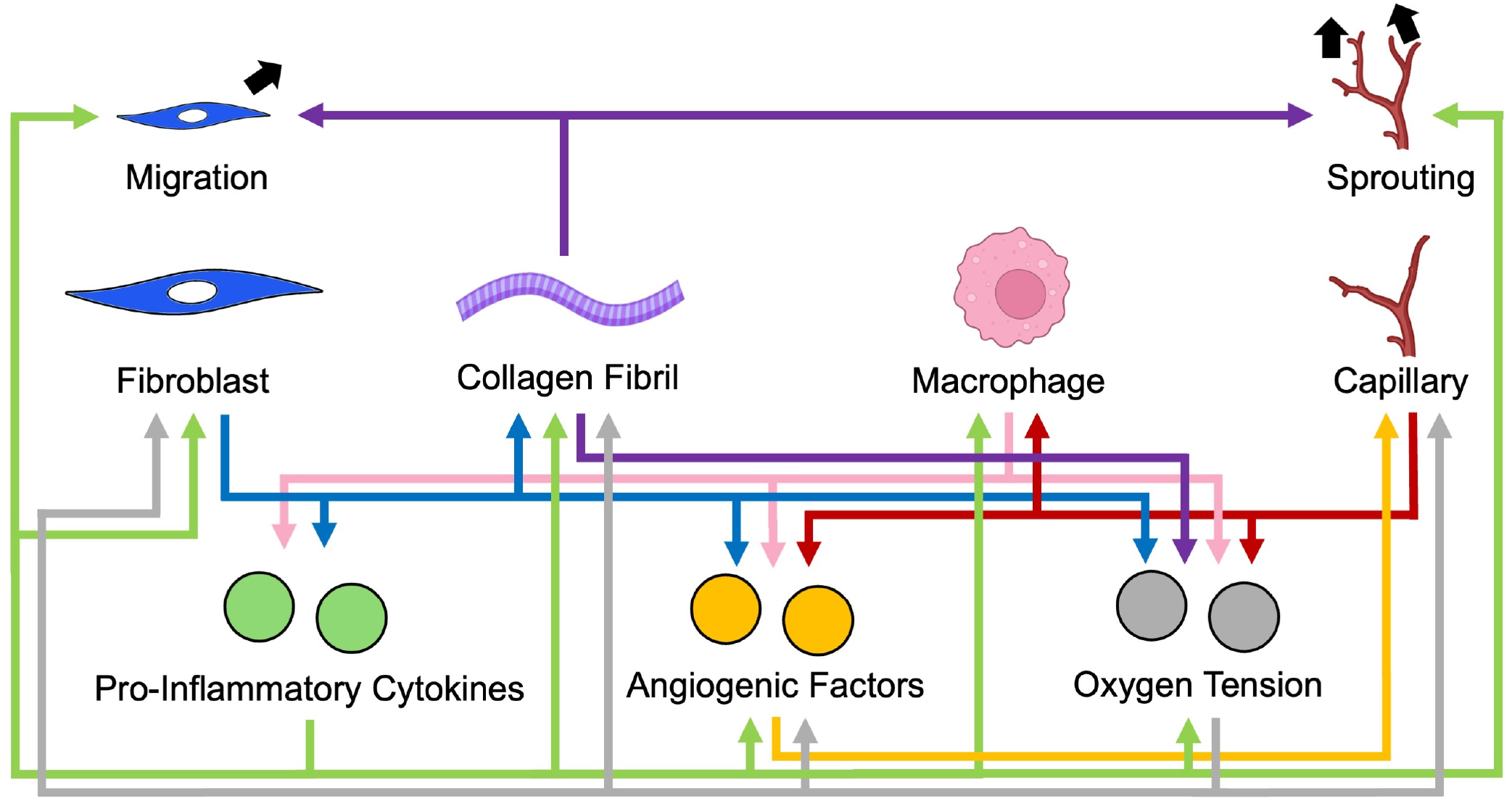
Coupling Structure of the Cavity Healing Model. Interaction network illustrating couplings among the modeled species in the cavity healing model. Arrows denote dependencies between species, representing interactions encoded through the coupled partial and ordinary differential equations governing transport, signaling, and regulatory processes. The network highlights the interplay among cellular activity, biochemical signaling, oxygen transport, and collagen deposition that governs cavity healing.

The geometry employed in the model is illustrated in Figure 3 and represents a generalized porcine breast derived from the preclinical lumpectomy study described by Puls et al. (2021)^13^, consistent with prior model calibration^29^. The lumpectomy cavity is represented as an ellipsoidal void (*a* = *b* = 1.5 cm, *c* = 0.6 cm) with a depth of 1.15 cm measured from the breast surface. The surrounding breast tissue is modeled as a half-ellipsoidal domain (*a* = *b* = 2.32 cm, *c* = 2 cm), corresponding to a quadrantectomy involving approximately one quarter of the total breast volume, and embedded within a rectangular region of connective tissue measuring 15 *×* 15 *×* 2 cm^3^.

**Figure 3:**
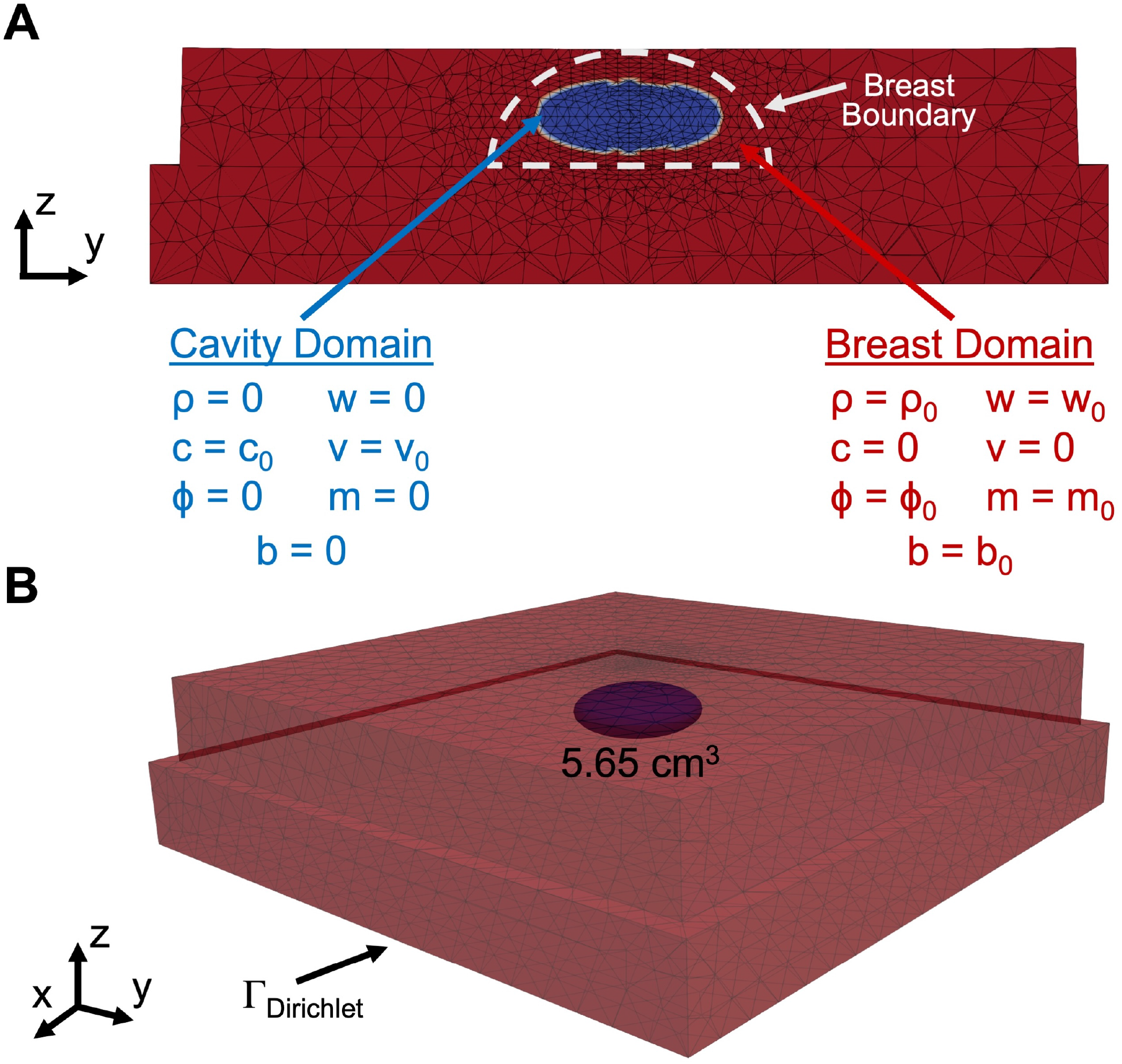
Porcine Breast Model Geometry and Domain Configuration. Finite element geometry and initial and boundary conditions for the porcine breast model shown in (A) cross-sectional and (B) isometric views. Based on the porcine lumpectomy model, the surgical cavity was assumed to be an ellipsoid with a volume of 5.65 cm^3^, while the surrounding breast domain was modeled as a half-ellipsoid (22.60 cm^3^), representing breast tissue geometry following quadrantectomy. Tissue external to the breast was modeled as connective tissue. The finite element mesh consisted of 97,517 tetrahedral elements. Initial conditions for fibroblast density (*ρ*), macrophage density (m), capillary density (b), pro-inflammatory cytokine concentration (c), angiogenic factor concentration (v), oxygen concentration (w), and collagen content (*ϕ*) are shown for the cavity and surrounding tissue. A Dirichlet boundary condition was applied along the chest wall boundary, while the exterior breast surface was treated as a free boundary.

### Experimental Data to Inform Model Calibration

Experimental data were incorporated to calibrate the predicted spatiotemporal profiles. Capillary and macrophage densities were quantified from histological sections collected from the porcine lumpectomy model at 1, 4, and 16 weeks following surgery, along with samples from healthy adipose and fibroglandular tissue^13^. As illustrated in Figure 4(i), histological sections were analyzed by sampling multiple regions across the cavity domain to quantify percent CD31-positive area (%CD31+), representing endothelial-lined capillary structures, and CD11b-positive inflammatory cell counts, which were assumed to primarily represent the macrophage population within the cavity region. To maintain consistency with the volumetric formulation of the model, these surface-based measurements were converted to volumetric densities. Displayed in Figure 4(ii), this was achieved using a simulation-based matching approach in which capillary networks and macrophage distributions were generated within representative volume elements (RVEs), and simulated histological slices were matched to experimentally observed measurements^38^. Additional details of the histological analysis and volumetric density estimation procedure are provided in the STAR Methods.

**Figure 4:**
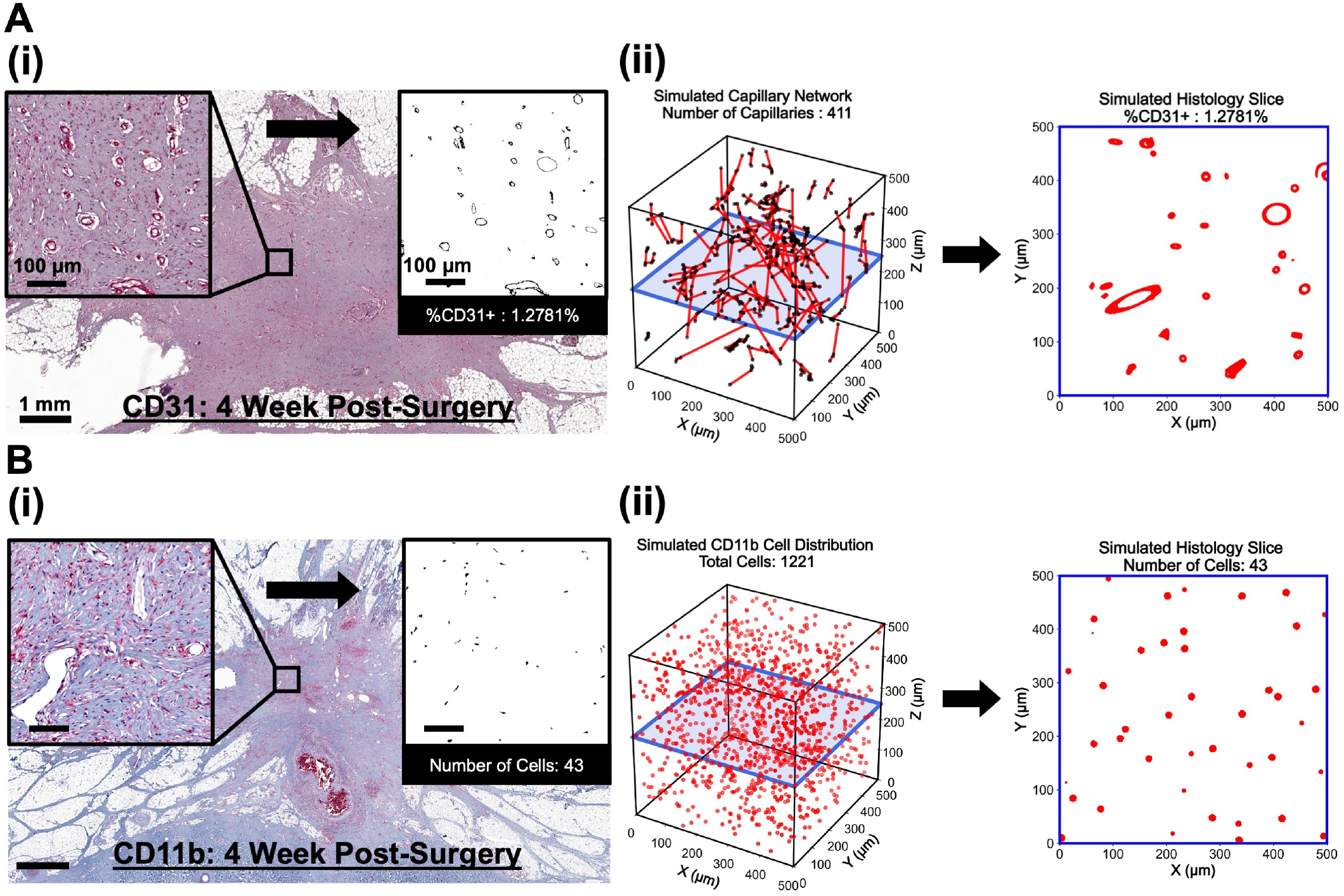
Histological Characterization of Capillary and Macrophage Distributions. Quantification of capillary area fraction and macrophage density from histological imaging, and estimation of corresponding volumetric densities for use in the computational model. (A) CD31 staining was used to visualize capillaries, while (B) CD11b staining was used to label macrophages in histological sections. (i) Histological images from porcine lumpectomy specimens (example shown at 4 weeks post-surgery) were subdivided into 500 *×* 500 *µm*^2^ regions of interest to quantify the percentage of CD31-positive area (%CD31+) as a measure of capillary area fraction and the number of CD11b-positive cells as a measure of macrophage density. (ii) To estimate volumetric densities, 500 *×* 500 *×* 500 *µm*^3^ representative volume elements were iteratively generated with simulated capillary networks or macrophage distributions until simulated histological slices matched the experimentally measured %CD31+ values or cell counts^38^.

Volumetric density estimates for macrophages and capillaries derived from histological measurements are summarized in Table 1 across both healthy breast tissue and postoperative time-points. For healthy tissue, volumetric densities were quantified separately for adipose and fibroglandular regions. In the model, a homogeneous tissue composition of approximately 70% adipose and 30% fibroglandular tissue was assumed^39^, yielding effective volumetric density values corresponding to the homogeneous breast tissue. Hematoma and/or seroma formation was observed both grossly and histologically at 1 week following lumpectomy, indicative of a cavity environment lacking vascularization and cellular infiltration. Accordingly, capillary and macrophage densities were assumed to be zero at this timepoint (Table 1). By 4 weeks, fibrovascular scar tissue was evident within the cavity, with capillary and macrophage densities elevated to approximately 2.0 and 2.4 times healthy tissue values, respectively (Table 1). At 16 weeks, fibrovascular scar tissue was maintained within the cavity, with capillary densities remaining elevated and macrophage densities showing a slight decline (Table 1).

**Table 1:**
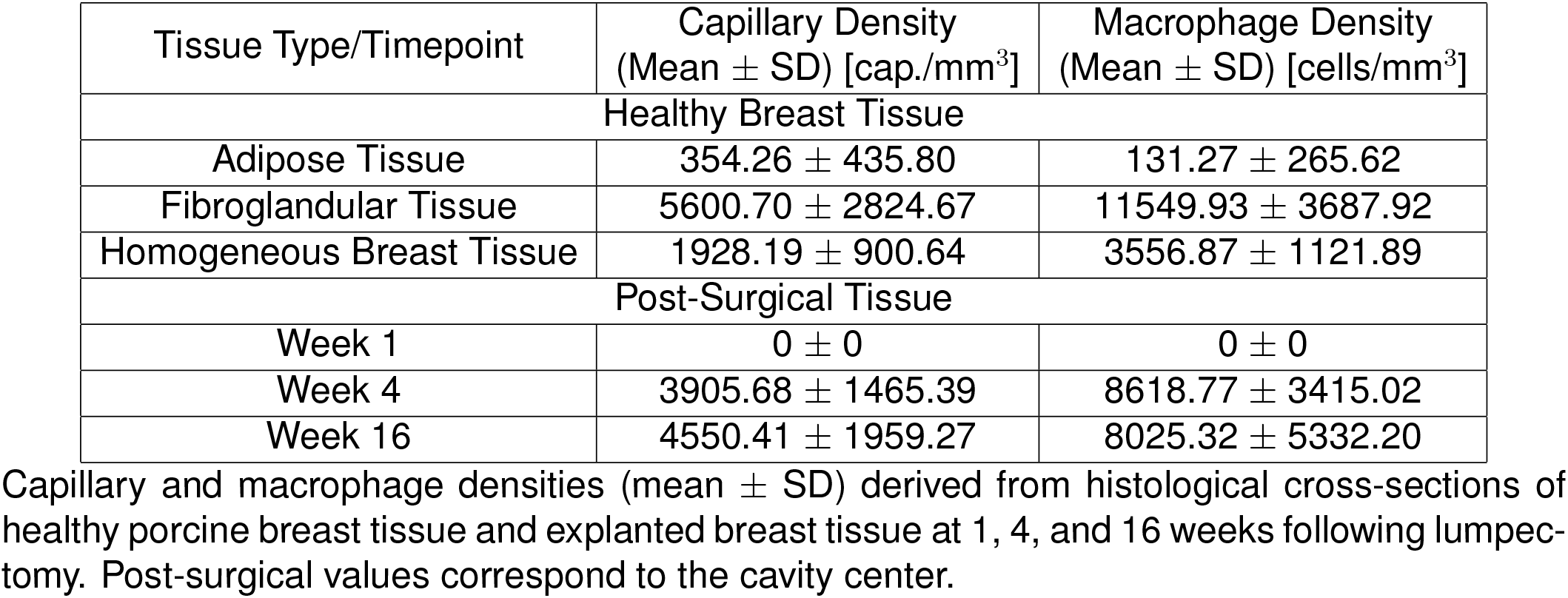
Histological Quantification of Capillary and Macrophage Densities. Capillary and macrophage densities (mean *±* SD) derived from histological cross-sections of healthy porcine breast tissue and explanted breast tissue at 1, 4, and 16 weeks following lumpectomy. Post-surgical values correspond to the cavity center.

| Tissue Type/Timepoint | Capillary Density<br>(Mean ± SD) [cap./mm<sup>3</sup>] | Macrophage Density<br>(Mean ± SD) [cells/mm<sup>3</sup>] |
| --- | --- | --- |
| Healthy Breast Tissue |  |  |
| Adipose Tissue | 354.26 ± 435.80 | 131.27 ± 265.62 |
| Fibroglandular Tissue | 5600.70 ± 2824.67 | 11549.93 ± 3687.92 |
| Homogeneous Breast Tissue | 1928.19 ± 900.64 | 3556.87 ± 1121.89 |
| Post-Surgical Tissue |  |  |
| Week 1 | 0 ± 0 | 0 ± 0 |
| Week 4 | 3905.68 ± 1465.39 | 8618.77 ± 3415.02 |
| Week 16 | 4550.41 ± 1959.27 | 8025.32 ± 5332.20 |

Beyond the histological measurements described above, additional experimental and literature-based data were incorporated to inform the modeled species. Fibroblast and collagen densities previously quantified from H&E-stained sections of the same porcine lumpectomy study are incorporated here as established measurements^29^. Pro-inflammatory cytokine and angiogenic factor levels were informed by clinical studies evaluating cytokine concentrations in post-surgical serous fluid collected from breast cancer patients. These data characterize their temporal behavior during healing, indicating that pro-inflammatory cytokine activity approximately resolves on the order of four weeks following surgery^40,41^. In contrast, angiogenic factor levels exhibit a transient increase, reaching a maximum of approximately 5 *±* 3.2 times the initial concentration within the cavity space^42^. Oxygen tension was informed by values reported in the literature, with early non-vascularized regions exhibiting low oxygen levels (0–5 mmHg)^43^, consistent with hematoma formation, while scar tissue maintains oxygen tensions slightly below those of healthy tissue^44,45^. In addition to these data sources, literature-informed relationships were used to guide the relative timing and progression of key biological processes with respect to one another. These include an early inflammatory response associated with macrophage activity preceding capillary and fibroblast responses^46–48^, as well as an earlier increase in angiogenic factor levels relative to capillary density^42,49^.

### Bayesian Calibration Using a Multi-Task Gaussian Process Surrogate

Within the computational model framework, several parameters governing the coupled biochemical system were informed through literature and experimental data, while 14 model parameters remained uninformed and could not be directly specified (Table 2). This provided an opportunity to calibrate these parameters by fitting model predictions to the histological and experimental data previously described. However, performing this calibration directly using the finite element framework is computationally prohibitive due to the high cost of evaluating the model across a large parameter space and the number of simulations required. To address this challenge, a multi-task Gaussian process (GP) surrogate model was employed to approximate the spatiotemporal response of the computational model. In contrast to a standard GP, which models each output independently, the multi-task formulation was implemented to account for correlations across biochemical species and over time, improving surrogate predictions for the coupled system^50^. As outlined in Figure 5, the surrogate model was constructed by first generating simulation data from the computational model across a broad range of parameter combinations defined in Table 2, which were then used to train the multi-task GP. The surrogate was subsequently refined through an iterative sampling strategy to improve predictive accuracy in underexplored regions of the parameter space.

**Table 2:** Parameters Optimized in the Computational Model. Uninformed model parameters included in the calibration procedure, with corresponding descriptions, prior ranges used for sampling, and the maximum likelihood estimate (MLE) obtained from Bayesian inference.

| Parameter | Description | Range | MLE Estimate |
| --- | --- | --- | --- |
| $p_{\rho,c}$ (1/hr) | Cytokine-Increased Fibroblast Proliferation | [0.015, 0.020] | 0.0166 |
| $p_{\phi,c}$ (1/hr) | Collagen Production Activated by Cytokine | $[5 \times 10^{-8}, 3.633 \times 10^{-7}]$ | $1.42 \times 10^{-7}$ |
| $d_w$ (1/hr) | Natural Oxygen Degradation | [0.001, 0.07] | 0.0501 |
| $p_{v,\rho}$ (1/hr) | Fibroblast Secretion of Angiogenic Factor | $[5 \times 10^{-13}, 1 \times 10^{-11}]$ | $2.44 \times 10^{-12}$ |
| $K_{v,\rho}$ (-) | Secretion of Angiogenic Factor Saturation | $[3.5 \times 10^{-5}, 7 \times 10^{-5}]$ | $4.71 \times 10^{-5}$ |
| $p_{v,m}$ (1/hr) | Macrophage Secretion of Angiogenic Factor | $[1 \times 10^{-12}, 1 \times 10^{-10}]$ | $2.88 \times 10^{-11}$ |
| $d_{v,b}$ (mm<sup>3</sup>/cap. · hr) | Angiogenic Factor Capillary Absorption Rate | $[5 \times 10^{-7}, 7.5 \times 10^{-6}]$ | $4.09 \times 10^{-6}$ |
| $d_v$ (1/hr) | Angiogenic Factor Decay | $[5 \times 10^{-4}, 7.5 \times 10^{-2}]$ | $4.42 \times 10^{-3}$ |
| $p_m$ (1/hr) | Macrophage Production | [0.005, 0.1] | 0.0553 |
| $p_{m,c}$ (1/hr) | Cytokine-Increased Macrophage Production | $[0.5 \cdot p_m, 5 \cdot p_m]$ | $0.9808 \cdot p_m$ |
| $K_{m,b}$ (-) | Capillary-Dependent Macrophage Saturation | [100, 2000] | 348.3534 |
| $d_{b,\phi}$ (-) | Capillary Diffusion Scaling Constant | [62.679, 12472.907] | 3707.5592 |
| $p_b$ (1/hr) | Capillary Growth Rate | [0.005, 0.05] | 0.0088 |
| $K_{b,b}$ (-) | Capillary Growth Saturation | [9640, 19280] | 9853.9264 |

**Figure 5:**
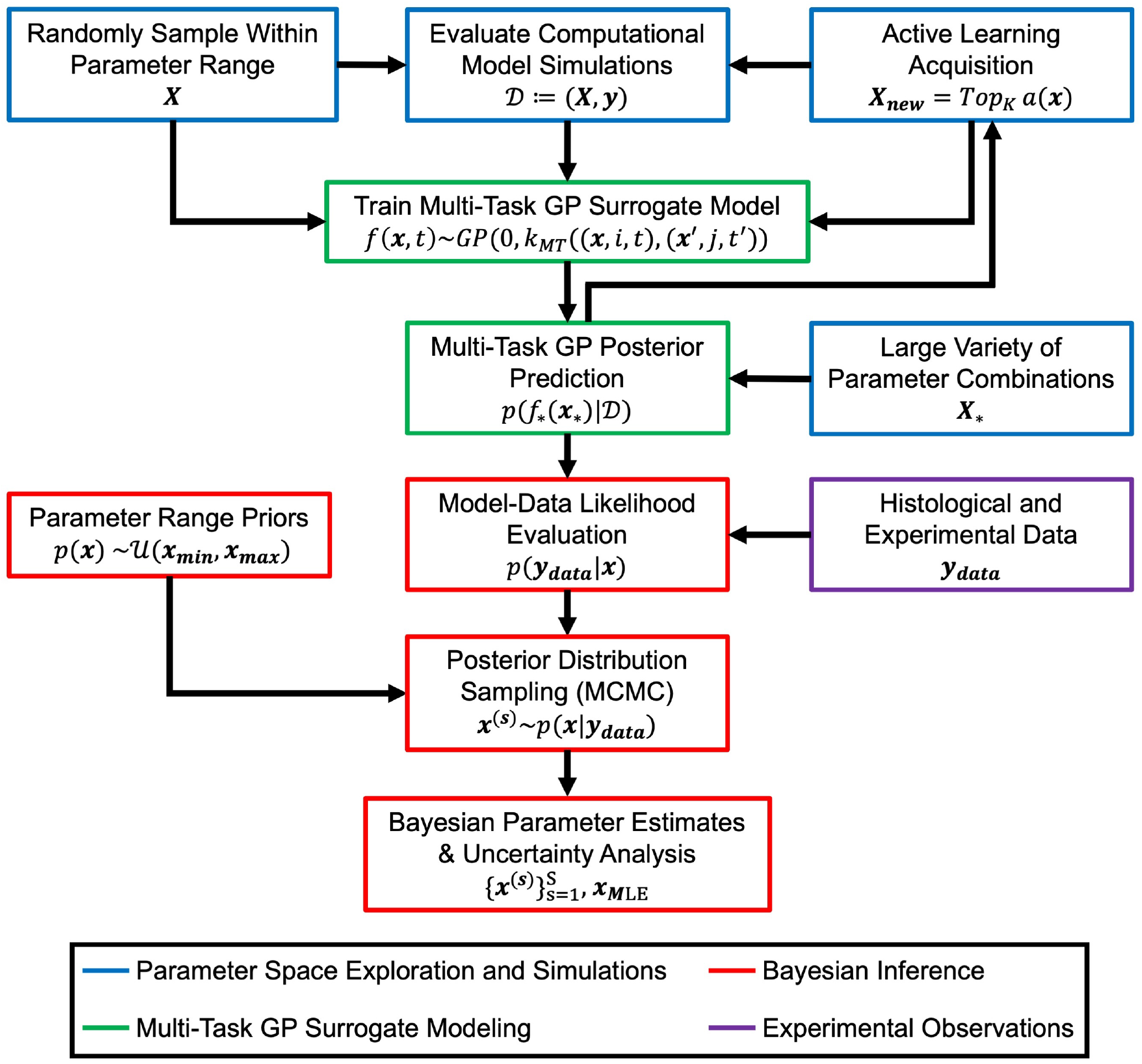
Surrogate Modeling and Bayesian Calibration Framework. Workflow integrating multi-task GP surrogate modeling and Bayesian calibration to align model predictions with experimental data through parameter optimization. Parameter combinations spanning the predefined parameter space are sampled and evaluated using the computational model to generate simulation data. These simulations are used to train a multi-task GP surrogate model that captures the temporal behavior of the modeled species. An active learning acquisition strategy identifies additional parameter combinations in regions of high predictive un-certainty, which are evaluated using the computational model and incorporated into the training data to iteratively improve surrogate accuracy. The trained GP surrogate is then used to evaluate likelihoods comparing model predictions with experimental data, and Bayesian inference is performed through MCMC sampling to obtain posterior parameter distributions that quantify uncertainty and characterize parameter values consistent with the experimental observations.

To evaluate the predictive accuracy of the surrogate model, repeated random train–test splits of the simulation dataset were performed over 25 iterations. In each iteration, the data were partitioned into an approximate 80% training and 20% testing split, where the multi-task GP was trained on the training subset and used to generate predictions for the held-out samples. These predictions were compared against the corresponding computational model simulations using the normalized root mean square error (NRMSE), computed for each species across time and normalized by the interquantile range of the training data. The surrogate demonstrated strong predictive accuracy, with mean NRMSE values of 0.046, 0.062, 0.082, 0.133, 0.111, and 0.086 for fibroblast density, collagen density, oxygen tension, angiogenic factor concentration, macrophage density, and capillary density respectively.

Once sufficiently trained, the GP surrogate enabled rapid and accurate evaluation of model responses across the parameter space, supporting efficient exploration of model behavior. This capability facilitates the implementation of a Bayesian calibration framework to estimate the uncertain model parameters. Unlike traditional calibration methods that seek a single optimal parameter set, the Bayesian approach represents parameter uncertainty through probability distributions, identifying a range of parameter combinations that are consistent with the experimental data. As illustrated in Figure 5, prior parameter ranges were defined and combined with likelihood-based comparison of model predictions and experimental observations to quantify agreement across the parameter space. The resulting posterior distribution was then explored using Markov Chain Monte Carlo (MCMC) sampling. These posteriors identify parameter combinations that best reproduce the experimental measurements while quantifying associated uncertainty in both the parameters and model predictions. Additional details regarding the GP surrogate formulation, training procedure, and Bayesian calibration framework are provided in the STAR Methods.

### Posterior Parameter Distributions and Identifiability

Within the Bayesian calibration framework, MCMC sampling yields posterior distributions of the model parameters that define the inferred parameter space. These distributions provide insight into the range of parameter values consistent with the experimental observations, as well as the degree of parameter constraint and interactions within the coupled system. The posterior parameter distributions are shown in Figure 6 as a series of corner plots, illustrating the marginal distributions and pairwise relationships across the 14 calibrated model parameters.

**Figure 6:**
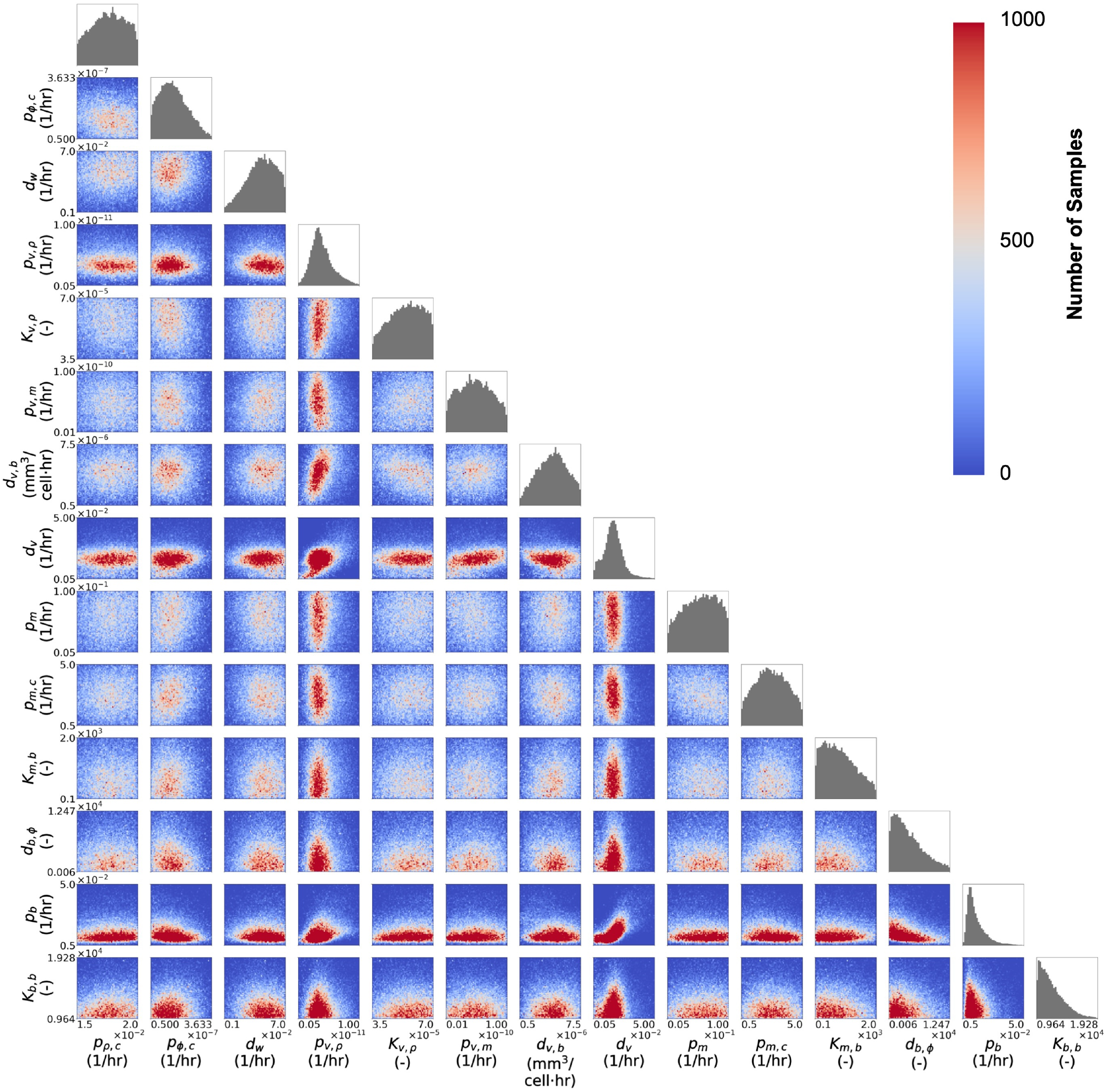
Posterior Parameter Distributions and Pairwise Relationships. Corner plot of posterior parameter samples obtained from Bayesian calibration of the cavity healing model using MCMC. Off-diagonal panels show pairwise relationships between model parameters, with the color mapping indicating posterior sample density and highlighting correlations between parameters. Diagonal panels show the marginal posterior distributions for each parameter.

Parameters associated with angiogenic factor and capillary processes are the most strongly identified, particularly those governing angiogenic factor secretion (*p*_*v,ρ*_) and loss (*d*_*v,b*_ and *d*_*v*_), as well as capillary transport (*d*_*b,ϕ*_) and growth (*p*_*b*_ and *K*_*b,b*_). These parameters exhibit narrow marginal distributions and well-defined pairwise relationships. This implies that the experimental data has direct information about some of these parameters, and there is strong coupling between angiogenic factor signaling and capillary formation (Figure 6). This behavior is consistent with the structure of the model, in which angiogenic factor directly regulates capillary growth. Oxygen behavior is also relatively well identified, as reflected in the relatively narrow posterior distribution of the oxygen degradation parameter (*d*_*w*_), reflecting its dependence on the evolving capillary network even though there is no direct measurement of oxygen in the dataset (Figure 6). Together, this hierarchy highlights the central role of vascularization and oxygen availability in governing the overall system response, with these tightly coupled processes exhibiting the strongest identifiability.

The parameters governing cytokine-increased fibroblast proliferation (*p*_*ρ,c*_) and macrophage production (*p*_*m*_, *p*_*m,c*_, and *K*_*m,b*_) are less well constrained, exhibiting broader posterior distributions and weaker pairwise structure (Figure 6). This reflects the fact that these cell populations are influenced by multiple upstream signals, including oxygen availability and inflammatory signaling. As a result, their effects can be compensated by other coupled mechanisms within the system, leading to increased flexibility in the parameter space. Despite their importance in co-ordinating tissue healing, this broader range of admissible values reflects reduced identifiability relative to more tightly coupled processes. The parameter governing collagen production activated by cytokine (*p*_*ϕ,c*_) is moderately constrained, with a posterior distribution that is less tightly defined than those governing vascularization (Figure 6). Even though collagen is the most down-stream variable, there is a tight coupling between collagen and capillary density, which explains why this downstream target is constrained by the data.

### Spatiotemporal Model Prediction of Healing Response

To evaluate the calibrated model behavior, the maximum likelihood estimate (MLE) parameter set was identified from the Bayesian calibration as the parameter combination that maximized the total log-likelihood across the MCMC samples. This parameter set represents the highest level of agreement between the model predictions and the experimental data and constraints used in the calibration. The corresponding MLE parameter values are provided in Table 2. When implemented within the computational model, this parameter set demonstrated strong agreement with the surrogate predictions, with NRMSE values below 0.04 for all species. This calibrated parameter set was then used to generate spatiotemporal predictions of the healing response, as shown in Figure 7.

**Figure 7:**
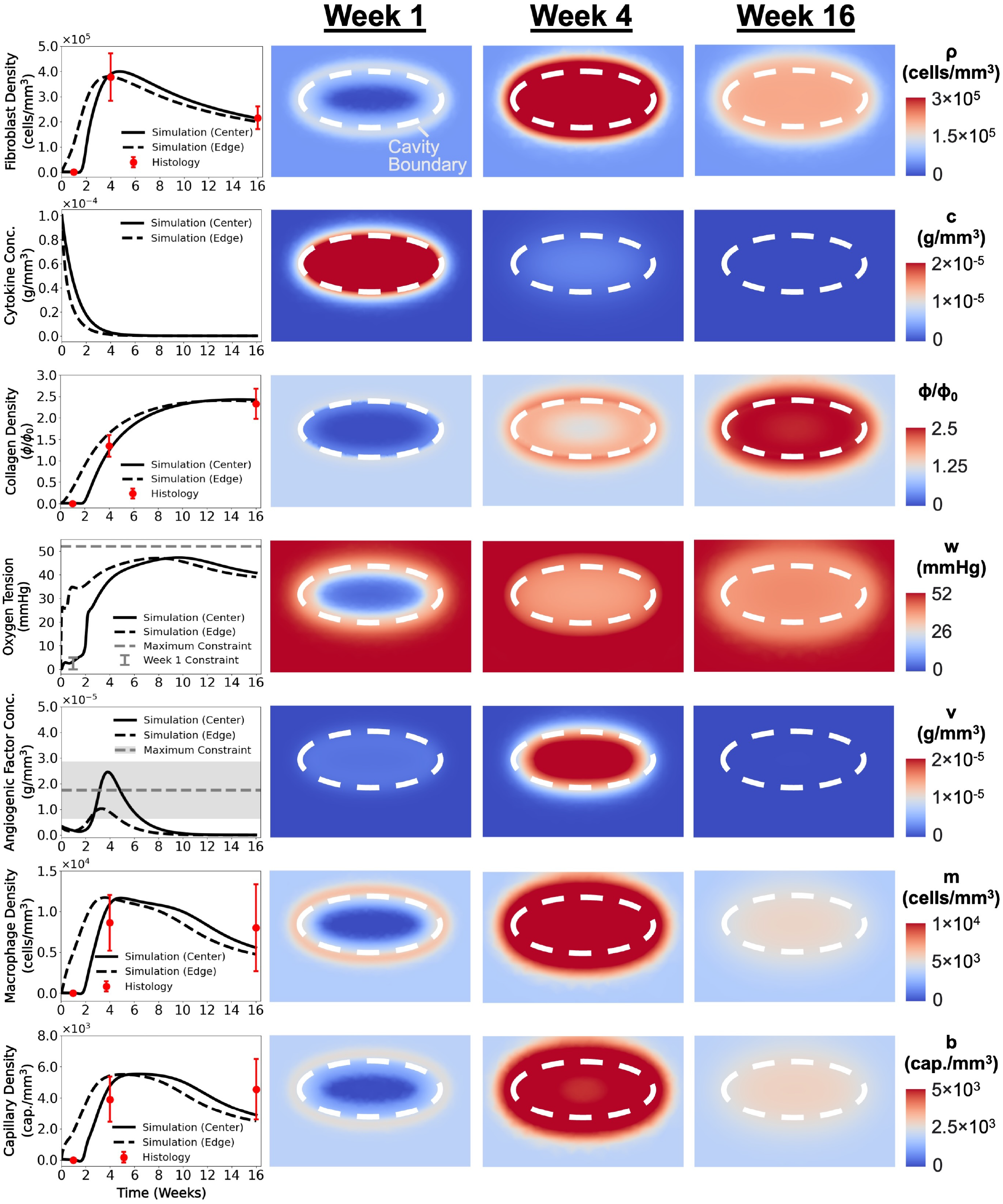
Spatiotemporal Distributions of Modeled Species During Cavity Healing. Simulation results of the cavity healing model using the optimized parameter set corresponding to the MLE identified through Bayesian calibration, representing the parameter set that maximizes agreement with the experimental data. Time-dependent trajectories of the modeled species are shown at the cavity center and cavity edge, alongside experimental histological measurements and imposed experimental constraints. Corresponding contour plots illustrate the spatial distributions of the modeled species at weeks 1, 4, and 16 following surgery.

In the early postoperative phase (week 1), the model captures the formation of a hematoma and/or seroma within the cavity center, resulting in negligible fibroblast, macrophage, capillary, and collagen densities in this region, consistent with histological observations (Figure 7). This lack of vascularization is accompanied by reduced oxygen tension within the cavity center, aligning with the imposed constraint of a hypoxic environment immediately following surgery. In contrast, pro-inflammatory cytokine levels remain elevated while declining from their initial peak, whereas angiogenic factor concentrations remain near their baseline values at this early time-point (Figure 7). Despite the lack of cellular and vascular presence at the cavity center, distinct spatial gradients are observed at the cavity boundary. Elevated macrophage density and early capillary presence occur along the edge of the cavity. Fibroblast and collagen densities also begin to increase along the cavity edge, though to a lesser extent than the more rapidly responding inflammatory and vascular components (Figure 7).

As the healing response progresses into the proliferative phase (week 4), the model captures a pronounced increase in fibroblast, macrophage, and capillary densities within the cavity (Figure 7). These populations are elevated relative to their corresponding healthy tissue levels and exhibit relatively uniform distributions between the cavity center and boundary, consistent with histological observations. Fibroblast proliferation and migration within the cavity space support progressive collagen deposition at this stage. Capillary sprouting and growth during this period enhance perfusion throughout the cavity. Pro-inflammatory cytokine concentration decays rapidly over the initial four-week period and approaches resolution by this timepoint, consistent with reported phases of wound healing^40,41^. In contrast, angiogenic factor levels increase over this period and reach a peak near week 4, aligning with the timing of angiogenic signaling observed in the literature^42^.

With the transition to the remodeling phase (week 16), fibroblast and macrophage densities decline from their respective peak values at week 4, while remaining similarly distributed between the cavity center and boundary and within experimentally observed ranges (Figure 7). Correspondingly, the rate of collagen deposition decreases as fibroblast number diminishes, with collagen density approaching a plateau. Capillary density remains elevated following its increase during the proliferative phase. Over this same period, angiogenic factor concentrations decline from their peak at week 4 and approach resolution, after which a gradual decrease in capillary density becomes evident (Figure 7). This behavior reflects the transition from active vascular growth to maturation and pruning of the vascular network. Oxygen tension remains elevated through this period, similar to levels observed at week 4, reflecting sustained perfusion within the remodeled tissue, though values remain below those of healthy tissue, consistent with observations of fibrovascular scar tissue^44,45^.

### Structure and Variability in Calibrated Modeled Responses

While the MLE parameter set provides a representative solution, the Bayesian calibration framework yields a distribution of parameter combinations that produce similarly strong agreement with the experimental data. To further examine this variability, posterior samples exhibiting the highest likelihood values were evaluated within the computational model, and the resulting simulations were used to represent the range of model responses consistent with the experimental data and imposed constraints. These simulations were then grouped using k-means clustering based on similarities in the temporal evolution of macrophage and capillary densities, enabling identification of distinct behaviors in the predicted healing response. These results are summarized in Figure 8, which illustrates the clustered temporal profiles of each species across the high-likelihood simulations, alongside their corresponding parameter distributions.

**Figure 8:**
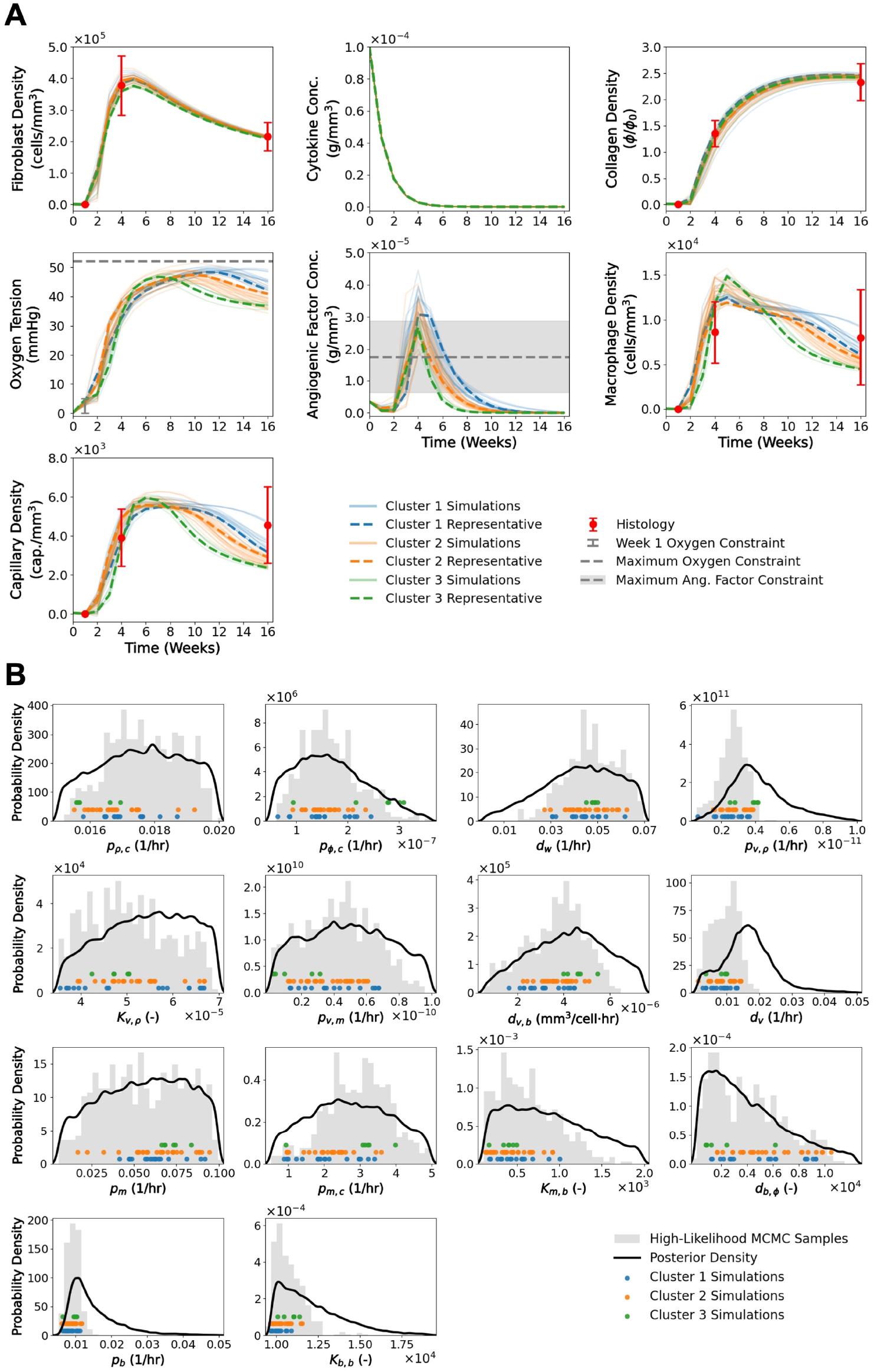
High-Likelihood Posterior Simulation Clusters and Associated Parameter Distributions. Clustering of the highest-likelihood posterior simulations obtained from Bayesian calibration of the cavity healing model. (A) Time-dependent profiles of model predictions at the cavity center for the 50 posterior simulations that maximize the log-likelihood, reflecting the best agreement between model predictions and experimental data. Simulations were grouped into three clusters using k-means clustering based on similarity in the temporal behavior of macrophages and capillaries. The representative simulation for each cluster corresponds to the parameter set with the highest likelihood within that cluster. (B) Marginal posterior distributions of model parameters inferred from Bayesian calibration, with markers indicating the parameter values corresponding to the clustered simulations. Gray histograms represent the distribution of the 500 parameter sets with the highest log-likelihood obtained from MCMC sampling.

Within these clustered simulations, variability in the predicted healing behavior is primarily driven by differences in the vascular dynamics of the model, reflected in the magnitude and persistence of capillary density across clusters (Figure 8A). Simulations grouped in Clusters 1 and 2 exhibit a sustained vascular response, characterized by a plateau in capillary density following its peak and maintaining elevated levels into the remodeling phase, with Cluster 2 displaying a shorter plateau and a more pronounced decline compared to Cluster 1 (Figure 8A). In contrast, simulations in Cluster 3 demonstrate a more transient behavior, with capillary density peaking earlier and undergoing a sharper decline that gradually slows toward the later remodeling phase (Figure 8A). These differences can be attributed to the corresponding angiogenic factor concentration, where more prolonged capillary responses correspond to higher levels and extended persistence of angiogenic signaling, while less sustained responses exhibit lower levels and a more rapid decline (Figure 8A). In relation to capillary density, oxygen tension reflects differences in perfusion, with more sustained vascularization corresponding to higher oxygen levels and reduced capillary presence associated with lower oxygen tension during the remodeling phase (Figure 8A). Consistent with these trends, macrophage density reaches a maximum at approximately week 4 in Clusters 1 and 2 and declines thereafter, with Cluster 1 remaining elevated longer into the remodeling phase, while Cluster 3 exhibits a delayed and higher maximum followed by a more rapid initial decline that gradually levels off at later timepoints (Figure 8A). In contrast to the variability observed in the vascular and inflammatory species, fibroblast and collagen densities exhibit reduced separation between clusters, with trajectories converging toward a similar range across simulations (Figure 8A), indicating diminished cluster-specific behavior.

Further insight into these cluster-dependent behaviors is provided by the corresponding parameter distributions (Figure 8B). Across both the high-likelihood MCMC samples and the clustered simulations, parameter values largely overlap within the posterior distributions, indicating that multiple parameter combinations can produce computational model simulations that match the experimental data and imposed constraints (Figure 8B). Within this overlapping parameter space, differences in parameter values are observed across clusters, indicating coordinated variation in processes governing angiogenesis. For example, simulations within Cluster 3 tend to exhibit a higher angiogenic factor capillary absorption rate (*d*_*v,b*_) alongside reduced macrophage secretion of angiogenic factors (*p*_*v,m*_), resulting in the observed lower angiogenic signaling and a diminished vascular response (Figure 8B). This behavior is further reinforced by a reduced capillary diffusion scaling constant (*d*_*b,ϕ*_), which reduces the rate of capillary growth throughout the cavity (Figure 8B). Increased macrophage production (*p*_*m*_ and *p*_*m,c*_) is also observed in simulations across Cluster 3, consistent with a compensatory response to the less sustained capillary presence, as macrophage supply is coupled to the vascular network in the model (Figure 8B). These patterns indicate that differences in healing trajectories arise from coordinated trade-offs among multiple coupled parameters, allowing consistent system-level behavior to emerge from a range of parameter combinations within the posterior distribution.

## DISCUSSION

Predictive models that can anticipate patient-specific healing trajectories and the physical changes underlying cosmetic outcomes following BCS have the potential to transform surgical planning by helping surgeons and patients balance oncologic, surgical, and cosmetic considerations. By evaluating treatment strategies before surgery, such models could guide shared patient-surgeon decision-making and support realistic expectations for postoperative recovery. This capability is clinically important because approximately one-third of patients experience post-surgical deformities such as indentations and breast asymmetry^33,51–53^, which can negatively impact patient satisfaction, quality of life, and recovery^6^. This variability reflects the patient-specific nature of the healing response, which remains difficult to predict and incorporate into clinical decision-making. To address this challenge, our prior work established a computational framework aimed at predicting healing and deformation following BCS to support personalized treatment planning^29,30^. This framework captured multiscale cell and tissue mechanics through the coupling of fibroblast activity, pro-inflammatory cytokine signaling, and collagen deposition. However, the model did not explicitly include inflammatory and vascular processes involved in the healing process. Many clinically relevant factors act through these pathways, linking patient-specific and treatment-related factors to variability in post-surgical healing. For instance, adjuvant radiation therapy can perturb the healing response following surgery, ultimately promoting fibrosis and altered tissue remodeling^33–35,54,55^. Similarly, diabetes, which affects approximately one-tenth of patients^56^, is associated with ischemic wound healing characterized by impaired vascularization and prolonged tissue repair^36,37^. Accordingly, incorporating inflammatory and vascular processes into the model framework is essential for characterizing the mechanobiological drivers underlying healing variability and advancing future patient-specific prediction of surgical outcomes following lumpectomy.

Over the past several decades, a number of mathematical and computational models have been developed to describe wound healing processes, with particular focus on angiogenesis^15–25^. Extending these concepts to post-BCS healing, Vavourakis et al. (2016) modeled cavity healing by applying cutaneous wound healing formulations^19,23,57^ within patient-specific breast geometries^28^. Thus, most prior models were developed for cutaneous wound healing or relied on idealized formulations, limiting their ability to capture experimental data on inflammatory and vascular remodeling within the unique case of cavity healing following BCS.

Building on these prior efforts, the present work develops a more detailed and physiologically representative formulation of the biochemical processes governing cavity healing. The current framework introduces a system of equations that captures the interactions between inflammatory signaling, fibroblast behavior, collagen deposition, angiogenic activity, and oxygen dynamics. These coupling relationships are informed by established physiological trends observed in experimental studies and prior modeling efforts^15,18,20,21,24,25^, ensuring that the interactions reflect known regulatory mechanisms underlying wound healing. A key advance of this work is the integration of experimental data to calibrate the model. To enable direct integration of the histological measurements, we further developed approaches for translating 2D histological data into volumetric quantities compatible with the wound healing simulations^38^. For instance, quantification of capillary density from %CD31+ histology measurements required the construction of capillary networks within RVEs^38^. The geometry and structure of these networks were informed by literature and validated against vascular architecture obtained from micro-computed tomography (microCT) imaging^49,58^. In summary, the present formulation integrates prior wound healing models, literature-informed physiological constraints, and porcine lumpectomy experimental data, resulting in a calibrated mechanistic framework for investigating coupled inflammatory and vascular remodeling during cavity healing.

This study relies on a machine learning surrogate model to enable Bayesian calibration, as direct calibration using the full finite element framework is computationally infeasible^59^. The surrogate model was trained on simulations from the computational framework to approximate the model response across the parameter space, reducing evaluation times from approximately 15 minutes per simulation to sub-second predictions. Gaussian process regression was employed due to its ability to provide predictive uncertainty alongside model outputs^60^. Specifically, a multi-task GP formulation was implemented to account for the coupled behavior across the modeled biochemical species, enabling shared information across outputs and improving predictive accuracy relative to standard single-task approaches^50^. This formulation further enabled the surrogate to capture both species-specific temporal dynamics and interactions among the coupled biological processes^61,62^. Within the Bayesian calibration framework, this approach enabled efficient exploration of parameter uncertainty^59^. The resulting posterior distributions revealed that multiple parameter combinations can reproduce experimentally observed healing trends, emphasizing that cavity healing is governed by coordinated interactions among coupled inflammatory, vascular, and ECM remodeling processes rather than a single unique parameter set (Figure 8).

The coupling between angiogenic signaling, capillary growth, and oxygen delivery is a central feature of the model and is reflected in the posterior parameter distributions (Figure 6), which show that parameters governing angiogenesis are the most tightly constrained. This suggests that the vascular response serves as a key regulator of cavity healing. Variability in angiogenic signaling and vascular development is the main contributor to qualitatively different model responses (Figure 8A), where healing trajectories are clustered based on variations in the magnitude and persistence of angiogenic factor expression and capillary density. Specifically, simulations exhibiting elevated and sustained capillary density correspond to increased oxygen tension and higher macrophage density, whereas reduced or more transient vascular responses are associated with lower oxygen levels and diminished macrophage presence (Figure 8A). Collectively, these results suggest that variability in angiogenic signaling and vascular remodeling may represent a major mechanistic source of heterogeneity in healing trajectories through its downstream effects on vascularization, oxygen delivery, and inflammatory activity.

In contrast to the vascular processes, the parameters governing collagen deposition are less tightly constrained in the posterior distributions (Figure 6), indicating a broader range of admissible values consistent with the experimental observations. This reflects the role of collagen as a downstream component that is nevertheless somewhat coupled to angiogenesis^63^. The posterior distributions associated with the cellular processes, namely fibroblast proliferation and macrophage production, were the least constrained (Figure 6). These cellular processes largely act downstream of the vascular environment.

As an initial extension of the computational framework to incorporate inflammatory and vascular processes, this study is not without limitations. First, this work does not incorporate the mechanobiological components included in prior formulations of the framework^29,30^. Here, the focus was placed on incorporating and calibrating inflammatory and vascular processes; therefore, mechanical contributions related to cavity contraction and deformation were not included. Future work will reintegrate these mechanobiological couplings to enable a fully coupled description of healing and deformation following BCS. Second, the current framework does not account for important patient-specific risk factors, such as age, diabetes, and smoking status, or adjuvant treatments including radiation therapy, all of which may influence vascular function and healing trajectories. While the present study focuses on establishing the underlying inflammatory and angiogenic dynamics, incorporating these factors represents an important step toward patient-specific predictions. Model calibration was based on available experimental data and literature, and future integration of longitudinal clinical measurements will be important for improving parameter constraints and strengthening predictive accuracy. Within the model, the vascular network is represented as a continuum capillary density field governed by simplified transport processes, rather than explicitly resolving discrete vessel structure or sprouting behavior. This formulation is appropriate for capturing the tissue-scale evolution of perfusion and oxygen delivery but may limit representation of localized heterogeneity in vascular architecture and transport. Finally, the proposed model represents a reduced biochemical system that focuses on key cell populations and signaling pathways to capture the dominant inflammatory and vascular processes of healing. Accordingly, specific inflammatory cell subtypes and individual cytokine signaling pathways are not explicitly resolved, with their effects represented through aggregate formulations. This simplification enables feasible calibration while preserving the primary features of the healing response. Expanding the model to include additional cell populations, signaling pathways, and vascular architecture would further refine its predictive capability and enable more detailed investigation of inflammatory and regenerative processes.

In conclusion, this study extends a computational framework for post-BCS cavity healing to incorporate experimentally calibrated inflammatory and vascular processes. Using histological data from a porcine lumpectomy model together with literature-informed physiological constraints, Bayesian calibration with a multi-task GP surrogate identified parameter ranges and regulatory relationships capable of reproducing observed healing trends. Through these simulations, the model revealed that vascularization plays a central regulatory role in cavity healing, with angiogenic signaling driving capillary development, oxygen restoration, cellular activity, and collagen deposition. Ultimately, this framework provides a foundation for patient-specific predictive tools that can evaluate post-surgical healing trajectories, and inform surgical planning and therapeutic strategies to improve cosmetic outcomes and quality of life following BCS.

## RESOURCE AVAILABILITY

### Lead contact

Requests for further information and resources should be directed to and will be fulfilled by the lead contact, Adrian Buganza Tepole.

### Materials availability

This study did not generate new materials.

### Data and code availability

- All data reported in this paper will be shared by the lead contact upon request.
- All original code generated in this study is publicly available at: https://github.com/zharbin/2026_BCS_AngModel.
- Any additional information required to reanalyze the data reported in this paper is available from the lead contact upon request.

## ACKNOWLEDGMENTS

The authors thank the Purdue University Histology Research Laboratory, a core facility of the NIH-funded Indiana Clinical and Translational Science Institute, and its staff for their assistance with histological slide preparation.

## AUTHOR CONTRIBUTIONS

Conceptualization, Z.H., H.G., S.V.-H., and A.B.T.; methodology, Z.H., R.A.M., H.G., S.V.-H., and A.B.T.; investigation, Z.H., H.G., S.V.-H., and A.B.T.; writing-–original draft, Z.H., H.G., S.V.-H., and A.B.T.; writing-–review & editing, Z.H., C.F., R.A.M., H.G., S.V.-H., and A.B.T.; funding and resources, Z.H., C.F., R.A.M., S.V.-H., and A.B.T.; supervision, C.F., H.G., S.V.-H., and A.B.T.

## SUPPLEMENTAL INFORMATION INDEX

Figure S1. Collagen-Dependent Capillary Diffusion Relationship

Table S1. Parameters for the Fibroblast Density Equation

Table S2. Parameters for the Pro-Inflammatory Cytokine Concentration Equation

Table S3. Parameters for the Collagen Density Equation

Table S4. Parameters for the Oxygen Tension Equation

Table S5. Parameters for the Angiogenic Factor Concentration Equation

Table S6. Parameters for the Macrophage Density Equation

Table S7. Parameters for the Capillary Density Equation

## STAR METHODS

### Key resources table

| REAGENT or RESOURCE | SOURCE | IDENTIFIER |
| --- | --- | --- |
| Antibodies |  |  |
| Recombinant Anti-CD31 Antibody | Abcam | Cat#ab182981; RRID: AB_2920881 |
| Recombinant Anti-CD11b Antibody | Abcam | Cat#ab133357; RRID: AB_2650514 |
| Horse Anti-Rabbit IgG ImmPress-AP Secondary Antibody | Vector Laboratories | Cat#MP-5401-15; RRID: AB_3148609 |
| Software and Algorithms |  |  |
| COMSOL Multiphysics | COMSOL AB | https://www.comsol.com/ |
| Aperio ImageScope | Leica Biosystems | https://www.leicabiosystems.com/us/digital-pathology/manage/aperio-imagescope/ |
| ImageJ | Schindelin et al.<sup>64</sup> | https://imagej.net/ij/ |
| Python version 3.9 | Python Software Foundation | https://www.python.org/ |
| GPyTorch | Gardner et al.<sup>65</sup> | https://gpytorch.ai/ |
| emcee | Foreman-Mackey et al.<sup>66</sup> | https://emcee.readthedocs.io/en/stable/ |
| Other |  |  |
| Negishi Community Cluster | Purdue Rosen Center for Advanced Computing | https://www.rcac.purdue.edu/compute/negishi |

## Method details

### Computational Model Formulation

The computational framework used in this study builds upon our previously developed mechanobiological models of wound healing, which were originally formulated to simulate cutaneous wound repair^67–69^ and later adapted to describe breast cavity healing following BCS^29,30^. In prior formulations, the biochemical component of the model accounted for fibroblasts, pro-inflammatory cytokines, and collagen. Here, the framework is extended to incorporate additional biochemical processes governing vascular remodeling and inflammatory signaling. The regulatory couplings introduced in this work are informed by experimental observations and prior computational models of wound healing and angiogenesis^15,18,20,21,24,25^, as illustrated in Figure 2. The resulting model is formulated as a coupled system of partial and ordinary differential equations and implemented using a finite element framework in COMSOL Multiphysics (COMSOL AB, Stockholm, Sweden). The governing equations for each modeled field are described below, with parameter descriptions and corresponding values provided in Tables S1–S7 in the Supplementary Material.

The evolution of fibroblast density (*ρ*) is described by

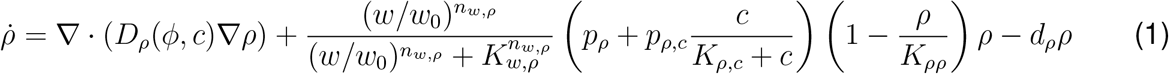

where the overdot denotes a time derivative. The source term describes fibroblast proliferation, which is enhanced in the presence of pro-inflammatory cytokines and modulated by local oxygen tension through an oxygen-dependent Hill function. The characteristic oxygen level defining the Hill response was informed by experimental observations suggesting reduced fibroblast proliferation below approximately 15 mmHg^70^. Fibroblast diffusion depends on both pro-inflammatory cytokine concentration and collagen density

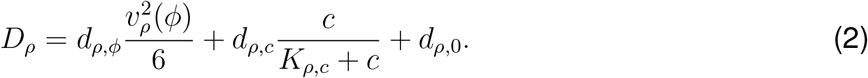

Fibroblast motility is modulated through the collagen-dependent fibroblast speed term *v*_*ρ*_(*ϕ*), which was informed by available in vivo wound healing data^71^. This diffusion formulation was originally developed and calibrated in our initial work describing the breast cavity healing mechanob ological model^29^.

Pro-inflammatory cytokines (c) are secreted by both fibroblast and macrophages

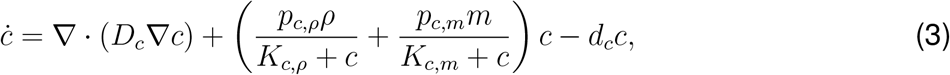

while cytokine concentration undergoes natural decay through the degradation term *d*_*c*_.

Collagen density (*ϕ*) reflects fibroblast-driven deposition enhanced by pro-inflammatory cytokines and regulated by local oxygen availability

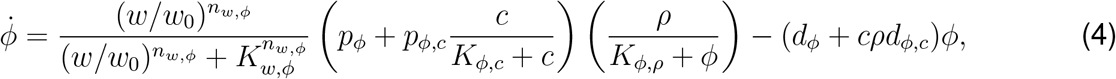

with the oxygen-dependent Hill function parameterized to reflect experimental observations indicating reduced collagen deposition under oxygen tensions below approximately 30–40 mmHg^72^

Oxygen tension (*w*) reflects supply from capillaries and consumption from both fibroblasts and macrophages

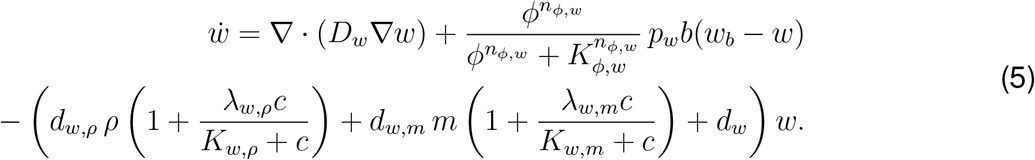

Here, *w*_*b*_ represents the oxygen tension at the arterial end of the capillaries (95 mmHg)^73^. Oxygen supply is modulated by a collagen-dependent Hill function, accounting for the immaturity of newly formed vessels in fibrin-rich early granulation tissue, which are less capable of delivering blood and oxygen than mature vessels^63,74^. Oxygen is consumed by fibroblasts and macrophages, which is enhanced in the presence of pro-inflammatory cytokines.

Similar to pro-inflammatory cytokines, angiogenic factors (*v*) are secreted by fibroblasts and macrophages, with production further enhanced in the presence of pro-inflammatory cytokines

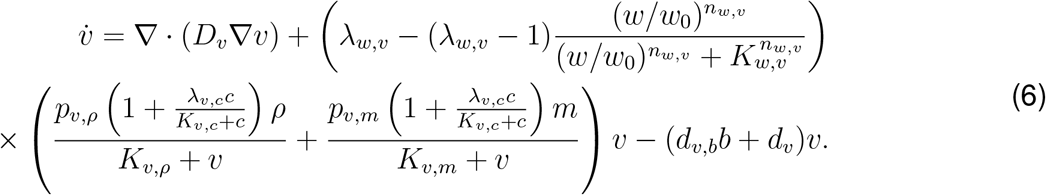

Production is also upregulated under hypoxic conditions through an oxygen-dependent response, resulting in an approximately threefold increase^75–78^. Angiogenic factors are also absorbed by capillaries and undergoes natural decay.

Macrophages (*m*) are supplied to the wound site through differentiation of monocytes migrating from the bloodstream, with recruitment regulated by capillary density and enhanced in the presence of pro-inflammatory cytokines

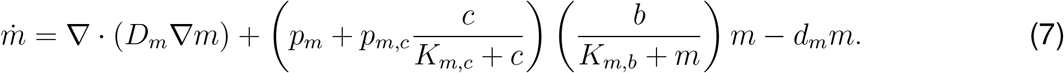

Capillary density (*b*) evolves through angiogenic growth driven by angiogenic factor concentration and regulated by local oxygen tension through an oxygen-dependent Hill response, using the same parameterization as that applied to collagen deposition

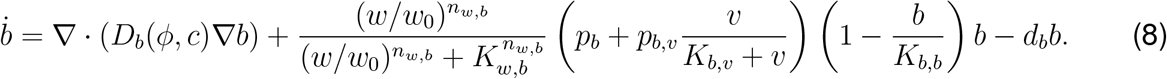

Capillary diffusion is described using the same formulation as fibroblast density, with dependencies on both collagen density and pro-inflammatory cytokine concentration

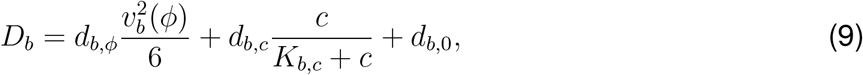

where motility is modulated through the collagen-dependent function *v*_*b*_(*ϕ*), informed by in vivo wound healing data^13,71^. The resulting *v*_*b*_(*ϕ*) relationship is shown in Figure S1.

### Histological Data Processing and Quantification

Histological samples were obtained from a preclinical porcine lumpectomy study^13^, and sections were immunohistochemically stained for CD31 and CD11b to identify capillaries and macrophages, respectively. Tissue explants were collected at 1, 4, and 16 weeks following surgery, as well as from healthy breast tissue. For each timepoint, 2–3 histological sections were analyzed, and within each section, 25 cross-sections measuring 500 *×* 500 *µm*^2^ were sampled spanning the cavity domain, as exemplified in Figure 4(i). These cross-sections were extracted using Aperio ImageScope (Leica Biosystems, Vista, CA, USA) and subsequently processed in ImageJ (National Institutes of Health, Bethesda, MD, USA) to quantify %CD31+ values and CD11b-positive cell counts. Image processing was performed primarily using color-based segmentation to isolate stained regions used to capture capillaries and macrophages^38^.

Histological surface measurements were converted to volumetric densities using a simulation-based matching approach (Figure 4(ii)) to align with the 3D finite element formulation of the computational model. Within this approach, RVEs were generated and populated with capillary networks (Figure 4A(ii)) and macrophage distributions (Figure 4B(ii)), from which simulated histological slices were extracted to reproduce the measured quantities^38^. For capillary quantification, three-dimensional capillary networks were generated within each RVE as branching structures, with branch order and segment lengths informed by literature and validated against vascular architectures^38,49,58^. Branch locations and orientations were assigned stochastically within the volume, with capillary cross-sectional dimensions informed by measurements obtained from porcine lumpectomy histological sections^38^. Virtual histological slices with a thickness of 4 *µm*, consistent with the experimental tissue sections, were extracted from the center of each RVE, and the corresponding %CD31+ value was computed based on the intersection of capillary segments with the slice plane^38^. Network generation was iteratively repeated until simulated values matched experimentally observed measurements, after which the corresponding volumetric capillary density was determined^38^. Macrophage quantification followed an analogous procedure, consistent with our prior work^29^. Cells of approximately 12 *µm* diameter, informed by histological observations and literature^79^, were distributed within the RVE. Virtual histological slices were then used to match simulated cell counts to experimentally observed values, enabling estimation of volumetric macrophage density.

### Multi-Task Gaussian Process Surrogate

Due to the computational expense of the finite element model, a multi-task Gaussian process surrogate was constructed to approximate the temporal response of the computational model across the biochemical species. The surrogate maps model parameters to outputs corresponding to multiple biochemical species evaluated at discrete timepoints at the center of the cavity, resulting in a multi-output regression problem in which each species–time pair is treated as a distinct task^61,62^. The model response is therefore represented as a function of the input parameters, species type, and time, denoted as *f*_*i*_(**x**, *t*). The multi-task Gaussian process is defined as

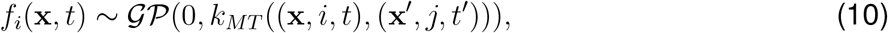

where *i* indicates the species type and **x** denotes the input parameters. A zero-mean prior is assumed and *k*_*MT*_ denotes the multi-task covariance kernel, which defines correlations between outputs across the input parameter space, species type, and time. To capture these dependencies, the kernel was constructed as the product of components corresponding to each of these dimensions

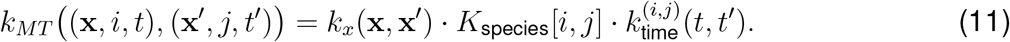

The input kernel *k*_*x*_(**x, x**^*′*^) was defined using a Rational Quadratic formulation with automatic relevance determination (ARD), allowing each model parameter to have an independent lengthscale. Inter-species dependencies were captured through a learned covariance matrix *K*_species_[*i, j*], parameterized via a Cholesky decomposition *K*_species_ = *LL*^*⊤*^, where *L* is lower triangular with positive diagonal entries. Temporal correlations were modeled using a convolutional squared exponential kernel with species-specific time lengthscales^62^, resulting in a time covariance that depends on the species pair

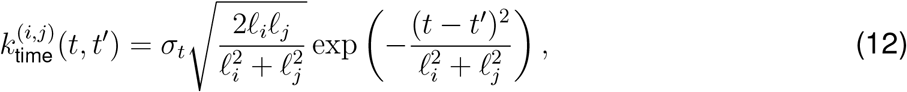

where *l*_*i*_ and *l*_*j*_ denote the time lengthscales associated with species *i* and *j*, respectively, and *σ*_*t*_ is a learned scaling parameter.

Training data for the surrogate were generated by sampling the uninformed model parameters using Latin hypercube sampling over the prescribed parameter ranges (Table 2), resulting in an initial dataset of 1,031 simulations. For each sampled parameter set, the computational model was evaluated, and model outputs were extracted at the center of the cavity for the biochemical species, excluding the pro-inflammatory cytokine as its temporal profile was not dependent on the uninformed parameters considered in the calibration. Each species response was evaluated at ten discrete timepoints spanning the 16-week healing period. Accordingly, each species–time pair was treated as a distinct task, resulting in a total of 60 tasks in the multi-task Gaussian process formulation. Model hyperparameters were learned by maximizing the marginal log-likelihood of the multi-task Gaussian process, implemented using GPyTorch^65^.

To improve surrogate accuracy in underexplored regions of the parameter space, the training dataset was also iteratively refined using an active learning strategy. Candidate parameter sets were generated using Latin hypercube sampling and evaluated based on a combined acquisition criterion that accounted for predictive uncertainty and coverage of the parameter space. Specifically, predictive variance from the multi-task GP and distance from existing training data were combined to rank candidate points. Top-ranked candidates were then refined using clustering and incorporated into the training dataset following evaluation with the computational model. This process was repeated over multiple iterations, resulting in an additional 485 simulations and progressively enhancing surrogate accuracy across the parameter space.

### Bayesian Calibration of Model Parameters

The trained surrogate model was subsequently used to perform Bayesian calibration of the uncertain model parameters, enabling efficient exploration of the parameter space while accounting for uncertainty in model predictions. Rather than identifying a single optimal parameter set, this approach seeks to characterize a distribution of parameter values that are consistent with the experimental observations through Bayesian inference. Let **y**_data_ denote the corresponding experimental observations. Following Bayes’ rule, the posterior distribution is given by

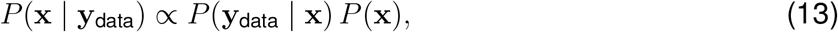

where *P* (**x** | **y**_data_) is the posterior distribution over the model parameters, *P* (**x**) represents the prior distribution, and *P* (**y**_data_ | **x**) is the likelihood function that quantifies agreement between model predictions and experimental observations. Independent uniform prior distributions were assigned to each parameter, with bounds corresponding to the ranges used in the initial Latin hypercube sampling (Table 2). The likelihood function was constructed from multiple components reflecting the different forms of data and constraints used for calibration

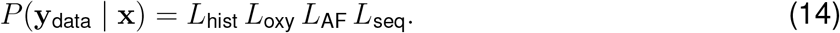

Species with direct histological measurements were incorporated through data-driven likelihood terms, while additional contributions were included to enforce physiologically informed constraints on oxygen tension, angiogenic factor behavior, and the relative timing of key biochemical processes.

For biochemical species with available histological measurements (i.e., fibroblast density, collagen density, macrophage density, and capillary density), likelihood terms were defined by comparing surrogate predictions to experimental observations at weeks 1, 4, and 16. Measurement error was modeled using Gaussian probability density functions (PDFs), with variance accounting for both experimental uncertainty and GP predictive variance^59^. Accordingly, the histology-based likelihood was defined as

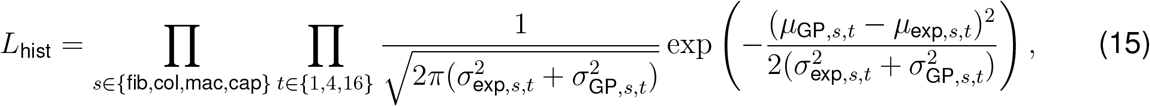

where *s* indexes the biochemical species and *t* denotes the observation timepoints. Here, *µ*_GP,*s,t*_ and 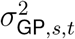 denote the GP-predicted mean and variance, while *µ*_exp,*s,t*_ and 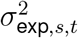 denote the corresponding histological measurements and uncertainties.

Oxygen tension was incorporated through literature-informed constraints reflecting early hypoxic conditions within the cavity (0–5 mmHg)^43^ and a late-stage upper bound consistent with oxygen levels in fibrovascular scar tissue, which remain lower than healthy breast tissue levels^44,45^. These constraints were implemented using Gaussian cumulative distribution functions (CDFs) to enforce physiologically relevant bounds on oxygen levels, such that

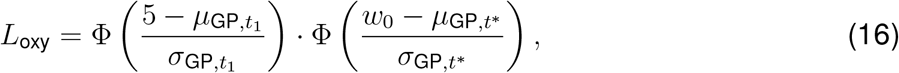

where Φ(·) denotes the standard normal cumulative distribution function, *t*_1_ corresponds to week 1, *t*^*^ is the timepoint at which the predicted oxygen tension attains its maximum value, and *w*_0_ represents the oxygen tension in healthy breast tissue (52 mmHg)^80^.

The angiogenic factor was incorporated through a constraint on its transient increase, reflecting literature-reported behavior in which concentrations rise to approximately 5 *±* 3.2 times the initial level during healing^42^. This behavior was enforced using a Gaussian PDF applied to the log-transformed fold-change, such that

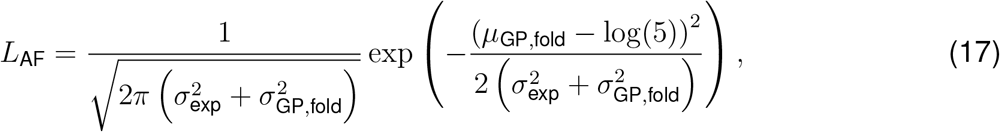

where *µ*_GP,fold_ denotes the GP-predicted log-transformed maximum fold-change, and 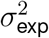 and 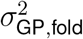 represent the corresponding experimental and GP predictive variances in log space.

Additional constraints were imposed on the relative timing of key biochemical processes to reflect expected cavity healing progression, including an early macrophage-mediated inflammatory response preceding fibroblast and capillary activity^46–48^, as well as an earlier increase in angiogenic factor relative to capillary density^42,49^. These relationships were enforced using Gaussian CDFs, such that

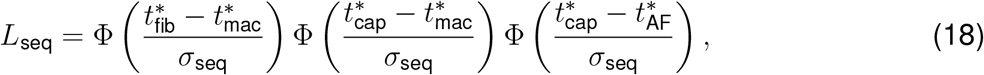

where *t*^*^ denotes the timepoint at which each species attains its maximum value, and *σ*_seq_ controls the softness of the timing constraints.

The posterior distribution was explored using MCMC sampling, with the trained GP surrogate used to efficiently evaluate the likelihood across the parameter space. Sampling was performed using an ensemble MCMC sampler implemented using emcee^66^, with 56 walkers used to explore the 14-dimensional parameter space. An initial burn-in phase of 2,000 steps was performed to allow the chains to converge toward high-probability regions, after which the sampler was reset and production sampling was conducted. Convergence was assessed using the integrated autocorrelation time and sampling was terminated once the total number of samples, *N*, satisfied the criterion *N* ≥ 50 *τ*_max_, where *τ*_max_ denotes the maximum autocorrelation time across all parameters. In practice, this resulted in on the order of 10^4^ production samples per walker.

### Quantification and statistical analysis

Histological measurements of capillary density and macrophage density were quantified to support model calibration. Detailed descriptions of the histological analysis and image processing procedures are provided in the STAR Methods, with summarized results reported in Table 1. For each timepoint (1, 4, and 16 weeks post-surgery, as well as healthy tissue), 2–3 histological sections were analyzed per staining type. Within each section, *n* = 25 cross-sections were strategically sampled across the cavity domain to capture spatial variability within the wound region. Measurements from these sampled regions were used to compute the mean and standard deviation (SD) at each timepoint. For volumetric density estimates (Table 1), variability reflects contributions from both the experimental measurements and the simulation-based conversion procedure.

## Supplementary Material

**Figure S1:**
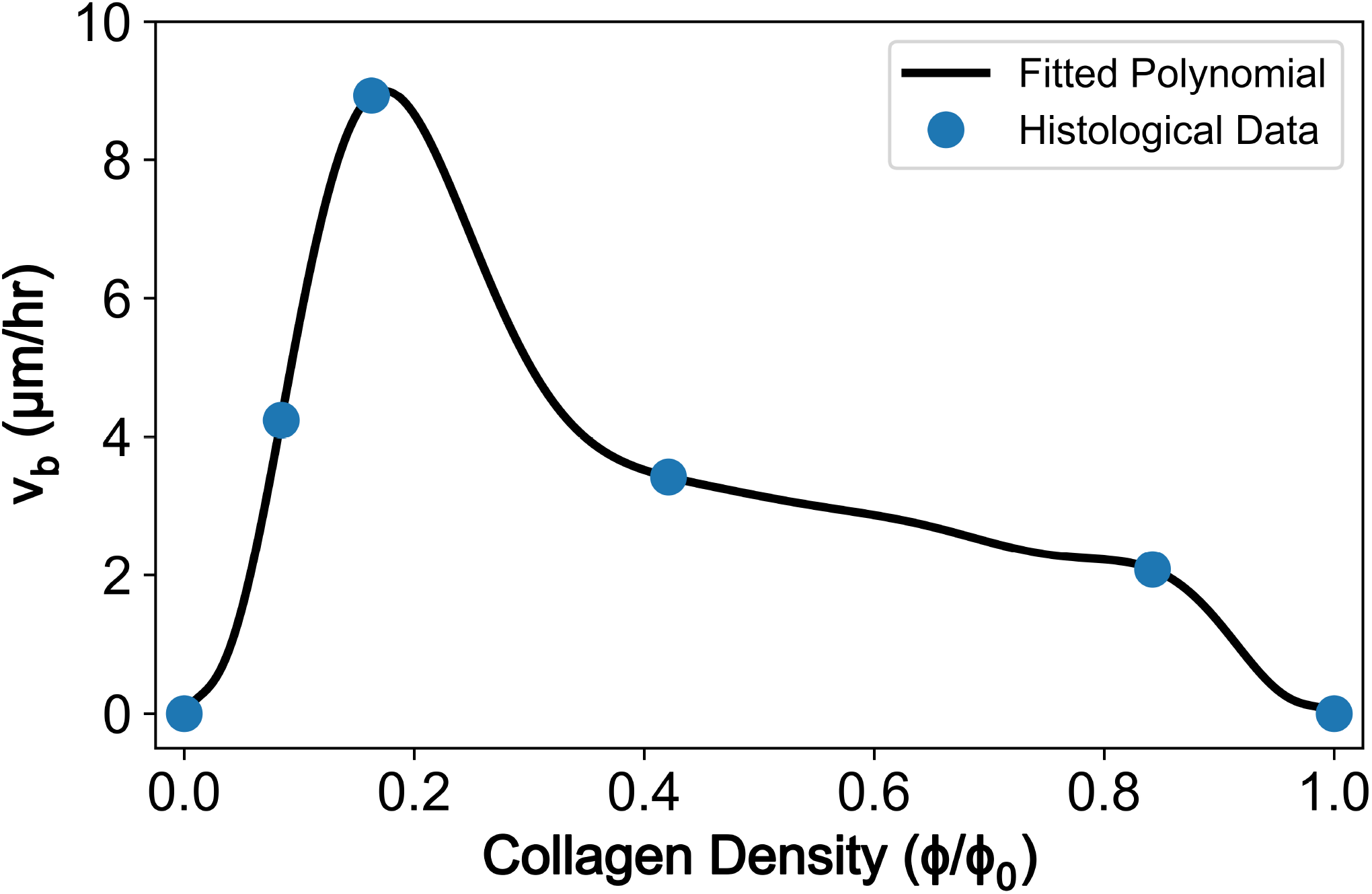
Collagen-Dependent Capillary Diffusion Relationship. Capillary growth speed with respect to collagen density (*v*_*b*_(*ϕ*)). The relationship was informed through analysis of histological images reported by [1, 2], where vascularization was evaluated within healing wounds treated with collagen scaffolds of varying collagen concentrations. Capillary growth speed was assumed to be zero at *ϕ* = 0 and *ϕ* = 1. A polynomial function was subsequently fit through the histological data points to define the collagen-dependent capillary diffusion behavior implemented within the computational model.

**Table S1:** Parameters for the Fibroblast Density Equation.

| **Term** | **Parameter Description** | **Value** | **Reference** |
| --- | --- | --- | --- |
| $\rho_0$ [cells/mm<sup>3</sup>] | Nominal Fibroblast Density | 55051 | [3] |
| $d_{\rho,\phi}$ [-] | Fibroblast Diffusion Scaling Constant | 1582.3 | [3] |
| $d_{\rho,c}$ [mm<sup>2</sup>/hr] | Cytokine-Increased Fibroblast Diffusivity | $6.12 \times 10^{-3}$ | [3] |
| $d_{\rho,0}$ [mm<sup>2</sup>/hr] | Baseline Fibroblast Diffusivity | $6.12 \times 10^{-5}$ | [3, 4] |
| $n_{w,\rho}$ [-] | Fibroblast Oxygen-Dependent Hill Function Exponent | 5 | Assumed based on [5] |
| $K_{w,\rho}$ [-] | Fibroblast Oxygen-Dependent Hill Function Saturation | 0.11 | Assumed based on [5] |
| $p_\rho$ [1/hr] | Fibroblast Proliferation | $9.00 \times 10^{-4}$ | [3] |
| $p_{\rho,c}$ [1/hr] | Cytokine-Increased Fibroblast Proliferation | 0.0166 | Calibrated in present study |
| $K_{\rho,c}$ [-] | Fibroblast Proliferation Saturation due to Cytokine | $1.00 \times 10^{-5}$ | [6] |
| $K_{\rho,\rho}$ [-] | Fibroblast Division Saturation | $10 * \rho_0$ | [7] |
| $d_\rho$ [1/hr] | Fibroblast Death Rate | $p_\rho(1 - \rho_0/K_{\rho,\rho})$ | [7] |

**Table S2:** Parameters for the Pro-Inflammatory Cytokine Concentration Equation.

| **Term** | **Parameter Description** | **Value** | **Reference** |
| --- | --- | --- | --- |
| $c_0$ [g/mm<sup>3</sup>] | Initial Cytokine Concentration Inside Cavity | $1.00 \times 10^{-4}$ | [6] |
| $D_c$ [mm<sup>2</sup>/hr] | Cytokine Diffusion Coefficient | 0.01208 | [8–10] |
| $p_{c,\rho}$ [1/hr] | Fibroblast Secretion of Cytokine | $1.635 \times 10^{-18}$ | [6] |
| $K_{c,\rho}$ [-] | Fibroblast Secretion of Cytokine Saturation | 1 | [6] |
| $p_{c,m}$ [1/hr] | Macrophage Secretion of Cytokine | $9.00 \times 10^{-17}$ | Assumed |
| $K_{c,m}$ [-] | Macrophage Secretion of Cytokine Saturation | $K_{\rho,c}$ | Assumed |
| $d_c$ [1/hr] | Cytokine Death Rate | 0.005 | [3] |

**Table S3:** Parameters for the Collagen Density Equation.

| **Term** | **Parameter Description** | **Value** | **Reference** |
| --- | --- | --- | --- |
| $n_{w,\phi}$ [-] | Collagen Oxygen-Dependent Hill Function Exponent | 4 | Assumed based on [11] |
| $K_{w,\phi}$ [-] | Collagen Oxygen-Dependent Hill Function Saturation | 0.1667 | Assumed based on [11] |
| $p_\phi$ [1/hr] | Collagen Production | $1.40 \times 10^{-8}$ | [3] |
| $p_{\phi,c}$ [1/hr] | Collagen Production Activated by Cytokine | $1.42 \times 10^{-7}$ | Calibrated in present study |
| $K_{\phi,c}$ [-] | Collagen Production Saturation due to Cytokine | $1.00 \times 10^{-4}$ | [6] |
| $K_{\phi,\rho}$ [-] | Collagen Production Saturation due to Collagen Fraction | 1.06 | [6] |
| $d_\phi$ [1/hr] | Collagen Degradation | $p_\phi(\rho_0/(K_{\phi,\rho} + 1))$ | [12] |
| $d_{\phi,c}$ [1/hr] | Collagen Degradation Activated by Cytokine | $8.81 \times 10^{-5}$ | [13] |

**Table S4:** Parameters for the Oxygen Tension Equation.

| **Term** | **Parameter Description** | **Value** | **Reference** |
| --- | --- | --- | --- |
| $w_0$ [mmHg] | Nominal Oxygen Tension | 52 | [14] |
| $D_w$ [mm<sup>2</sup>/hr] | Oxygen Diffusion Constant | 0.18 | [15] |
| $w_b$ [mmHg] | Oxygen Tension in Blood | 95 | [16] |
| $n_{\phi,w}$ [-] | Oxygen Collagen-Dependent Hill Function Exponent | 15 | Assumed based on [17] |
| $K_{\phi,w}$ [-] | Oxygen Collagen-Dependent Hill Function Saturation | 0.15 | Assumed based on [17] |
| $p_w$ [-] | Oxygen Production Term | $\frac{(d_{w,\rho}\rho_0 + d_{w,m}m_0 + d_w)w_0}{b_0(w_b - w_0)}$ | — |
| $d_{w,\rho}$ [mm<sup>3</sup>cell<sup>-1</sup>hr<sup>-1</sup>] | Oxygen Consumption by Fibroblasts | $9.60 \times 10^{-7}$ | [18] |
| $\lambda_{w,\rho}$ [-] | Fibroblast Oxygen Consumption Enhancement in Presence of Cytokines | $\lambda_{c,m}$ | Assumed |
| $K_{w,\rho}$ [-] | Fibroblast Oxygen Consumption Saturation due to Cytokines | $K_{\rho,c}$ | Assumed |
| $d_{w,m}$ [mm<sup>3</sup>cell<sup>-1</sup>hr<sup>-1</sup>] | Oxygen Consumption by Macrophages | $3.07 \times 10^{-6}$ | [19] |
| $\lambda_{c,m}$ [-] | Macrophage Oxygen Consumption Enhancement in Presence of Cytokines | 0.695 | [19] |
| $K_{c,m}$ [-] | Macrophage Oxygen Consumption Saturation due to Cytokines | $K_{\rho,c}$ | Assumed |
| $d_w$ [1/hr] | Natural Oxygen Degradation | 0.0501 | Calibrated in present study |

**Table S5:**
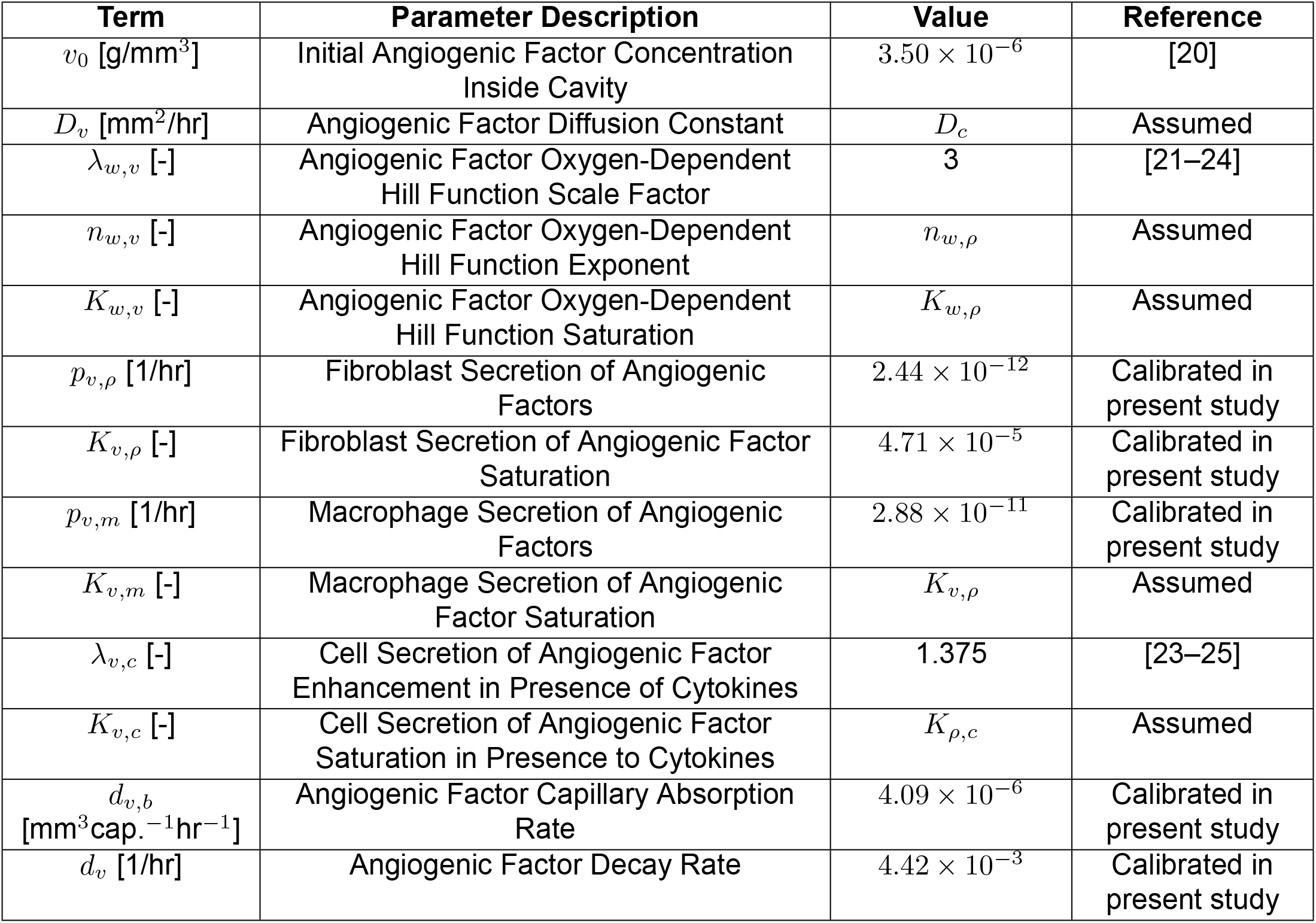
Parameters for the Angiogenic Factor Concentration Equation.

| **Term** | **Parameter Description** | **Value** | **Reference** |
| --- | --- | --- | --- |
| $v_0$ [g/mm<sup>3</sup>] | Initial Angiogenic Factor Concentration Inside Cavity | $3.50 \times 10^{-6}$ | [20] |
| $D_v$ [mm<sup>2</sup>/hr] | Angiogenic Factor Diffusion Constant | $D_c$ | Assumed |
| $\lambda_{w,v}$ [-] | Angiogenic Factor Oxygen-Dependent Hill Function Scale Factor | 3 | [21–24] |
| $n_{w,v}$ [-] | Angiogenic Factor Oxygen-Dependent Hill Function Exponent | $n_{w,\rho}$ | Assumed |
| $K_{w,v}$ [-] | Angiogenic Factor Oxygen-Dependent Hill Function Saturation | $K_{w,\rho}$ | Assumed |
| $p_{v,\rho}$ [1/hr] | Fibroblast Secretion of Angiogenic Factors | $2.44 \times 10^{-12}$ | Calibrated in present study |
| $K_{v,\rho}$ [-] | Fibroblast Secretion of Angiogenic Factor Saturation | $4.71 \times 10^{-5}$ | Calibrated in present study |
| $p_{v,m}$ [1/hr] | Macrophage Secretion of Angiogenic Factors | $2.88 \times 10^{-11}$ | Calibrated in present study |
| $K_{v,m}$ [-] | Macrophage Secretion of Angiogenic Factor Saturation | $K_{v,\rho}$ | Assumed |
| $\lambda_{v,c}$ [-] | Cell Secretion of Angiogenic Factor Enhancement in Presence of Cytokines | 1.375 | [23–25] |
| $K_{v,c}$ [-] | Cell Secretion of Angiogenic Factor Saturation in Presence to Cytokines | $K_{\rho,c}$ | Assumed |
| $d_{v,b}$ [mm<sup>3</sup>cap.<sup>-1</sup>hr<sup>-1</sup>] | Angiogenic Factor Capillary Absorption Rate | $4.09 \times 10^{-6}$ | Calibrated in present study |
| $d_v$ [1/hr] | Angiogenic Factor Decay Rate | $4.42 \times 10^{-3}$ | Calibrated in present study |

**Table S6:** Parameters for the Macrophage Density Equation.

| **Term** | **Parameter Description** | **Value** | **Reference** |
| --- | --- | --- | --- |
| $m_0$<br>[cells/mm<sup>3</sup>] | Nominal Macrophage Density | 3557 | Estimated in present study |
| $D_m$ [mm<sup>2</sup>/hr] | Macrophage Diffusion Constant | 0.005 | [26] |
| $p_m$ [1/hr] | Macrophage Production Term | 0.0553 | Calibrated in present study |
| $p_{m,c}$ [1/hr] | Cytokine-Increased Macrophage Production | $0.9808 \cdot p_m$ | Calibrated in present study |
| $K_{m,c}$ [-] | Macrophage Proliferation Saturation due to Cytokine | $K_{\rho,c}$ | Assumed |
| $K_{m,b}$ [-] | Capillary-Dependent Macrophage Saturation | 348.3534 | Calibrated in present study |
| $d_m$ [1/hr] | Macrophage Death Rate | $p_m(b_0/(K_{m,b}+m_0))$ | — |

**Table S7:** Figure S1: Collagen-Dependent Capillary Diffusion Relationship.

| **Term** | **Parameter Description** | **Value** | **Reference** |
| --- | --- | --- | --- |
| $b_0$ [cap./mm<sup>3</sup>] | Nominal Capillary Density | 1928 | Estimated in present study |
| $d_{b,\phi}$ [-] | Capillary Diffusion Scaling Constant | 3707.5592 | Calibrated in present study |
| $d_{b,c}$ [mm<sup>2</sup>/hr] | Cytokine-Increased Capillary Diffusivity | $d_{\rho,c}$ | Assumed |
| $K_{b,c}$ [-] | Cytokine-Increased Capillary Saturation Term | $K_{\rho,c}$ | Assumed |
| $d_{b,0}$ [mm<sup>2</sup>/hr] | Baseline Capillary Diffusivity | $d_{\rho,0}$ | Assumed |
| $n_{w,b}$ [-] | Capillary Oxygen-Dependent Hill Function Exponent | $n_{w,\phi}$ | Assumed based on [17] |
| $K_{w,b}$ [-] | Capillary Oxygen-Dependent Hill Function Saturation | $K_{w,\phi}$ | Assumed based on [17] |
| $p_b$ [1/hr] | Capillary Production Term | 0.0088 | Calibrated in present study |
| $p_{b,v}$ [1/hr] | Capillary Production in Presence of Angiogenic Factor | $0.88 \cdot p_b$ | [27] |
| $K_{b,v}$ [-] | Capillary Production Saturation due to Angiogenic Factor | $v_0/10$ | Assumed |
| $K_{b,b}$ [-] | Capillary Growth Saturation | 9853.9264 | Calibrated in present study |
| $d_b$ [1/hr] | Capillary Death Rate | $p_b(1 - b_0/K_{b,b})$ | — |

## References

1. Siegel, R.L., Kratzer, T.B., Wagle, N.S., Sung, H., and Jemal, A. (2026). Cancer statistics, 2026. Ca 76, e70043.

2. Wagle, N.S., Nogueira, L., Devasia, T.P., Mariotto, A.B., Yabroff, K.R., Islami, F., Jemal, A., Alteri, R., Ganz, P.A., and Siegel, R.L. (2025). Cancer treatment and survivorship statistics, 2025. CA: A cancer journal for clinicians 75, 308–340.

3. Fourquet, A., Campana, F., Zafrani, B., Mosseri, V., Vielh, P., Durand, J.C., and Vilcoq, J.R. (1989). Prognostic factors of breast recurrence in the conservative management of early breast cancer: a 25-year follow-up. International Journal of Radiation Oncology* Biology* Physics 17, 719–725.

4. Group, E.B.C.T.C. et al. (2005). Effects of radiotherapy and of differences in the extent of surgery for early breast cancer on local recurrence and 15-year survival: an overview of the randomised trials. The Lancet 366, 2087–2106.

5. Nelson, J.A., Rubenstein, R.N., Haglich, K., Chu, J.J., Yin, S., Stern, C.S., Morrow, M., Mehrara, B.J., Gemignani, M.L., and Matros, E. (2022). Analysis of a trend reversal in us lumpectomy rates from 2005 through 2017 using 3 nationwide data sets. JAMA surgery 157, 702–711.

6. de Boniface, J., Szulkin, R., and Johansson, A.L. (2021). Survival after breast conservation vs mastectomy adjusted for comorbidity and socioeconomic status: a swedish national 6-year follow-up of 48 986 women. JAMA Surgery 156, 628–637.

7. Chatterjee, A., Pyfer, B., Czerniecki, B., Rosenkranz, K., Tchou, J., and Fisher, C. (2015). Early postoperative outcomes in lumpectomy versus simple mastectomy. journal of Surgical Research 198, 143–148.

8. Litière, S., Werutsky, G., Fentiman, I.S., Rutgers, E., Christiaens, M.R., Van Limbergen, E., Baaijens, M.H., Bogaerts, J., and Bartelink, H. (2012). Breast conserving therapy versus mastectomy for stage i–ii breast cancer: 20 year follow-up of the eortc 10801 phase 3 randomised trial. The Lancet Oncology 13, 412–419.

9. Mogal, H.D., Clark, C., Dodson, R., Fino, N.F., and Howard-McNatt, M. (2017). Outcomes after mastectomy and lumpectomy in elderly patients with early-stage breast cancer. Annals of Surgical Oncology 24, 100–107.

10. Veronesi, U., Cascinelli, N., Mariani, L., Greco, M., Saccozzi, R., Luini, A., Aguilar, M., and Marubini, E. (2002). Twenty-year follow-up of a randomized study comparing breast-conserving surgery with radical mastectomy for early breast cancer. New England Journal of Medicine 347, 1227–1232.

11. Hau, E., Browne, L., Capp, A., Delaney, G.P., Fox, C., Kearsley, J.H., Millar, E., Nasser, E.H., Papadatos, G., and Graham, P.H. (2013). The impact of breast cosmetic and functional outcomes on quality of life: long-term results from the st. george and wollongong randomized breast boost trial. Breast Cancer Research and Treatment 139, 115–123.

12. Waljee, J.F., Hu, E.S., Ubel, P.A., Smith, D.M., Newman, L.A., and Alderman, A.K. (2008). Effect of esthetic outcome after breast-conserving surgery on psychosocial functioning and quality of life. Journal of Clinical Oncology 26, 3331–3337.

13. Puls, T.J., Fisher, C.S., Cox, A., Plantenga, J.M., McBride, E.L., Anderson, J.L., Goergen, C.J., Bible, M., Moller, T., and Voytik-Harbin, S.L. (2021). Regenerative tissue filler for breast conserving surgery and other soft tissue restoration and reconstruction needs. Scientific Reports 11, 2711.

14. Henderson, I.C. (2015). Breast cancer: Fundamentals of evidence-based disease management. Oxford University Press.

15. Pettet, G., Byrne, H., McElwain, D., and Norbury, J. (1996). A model of wound-healing angiogenesis in soft tissue. Mathematical biosciences 136, 35–63.

16. Olsen, L., Sherratt, J.A., Maini, P.K., and Arnold, F. (1997). A mathematical model for the capillary endothelial cell-extracellular matrix interactions in wound-healing angiogenesis. Mathematical Medicine and Biology: A Journal of the IMA 14, 261–281.

17. Byrne, H., Chaplain, M., Evans, D., and Hopkinson, I. (2000). Mathematical modelling of angiogenesis in wound healing: comparison of theory and experiment. Computational and Mathematical Methods in Medicine 2, 175–197.

18. Gaffney, E., Pugh, K., Maini, P., and Arnold, F. (2002). Investigating a simple model of cutaneous wound healing angiogenesis. Journal of mathematical biology 45, 337–374.

19. Maggelakis, S.A. (2003). A mathematical model of tissue replacement during epidermal wound healing. Applied Mathematical Modelling 27, 189–196.

20. Schugart, R.C., Friedman, A., Zhao, R., and Sen, C.K. (2008). Wound angiogenesis as a function of tissue oxygen tension: a mathematical model. Proceedings of the National Academy of Sciences 105, 2628–2633.

21. Xue, C., Friedman, A., and Sen, C.K. (2009). A mathematical model of ischemic cutaneous wounds. Proceedings of the National Academy of Sciences 106, 16782–16787.

22. Machado, M.J., Watson, M.G., Devlin, A.H., Chaplain, M.A., McDougall, S.R., and Mitchell, C.A. (2011). Dynamics of angiogenesis during wound healing: a coupled in vivo and in silico study. Microcirculation 18, 183–197.

23. Valero, C., Javierre, E., García-Aznar, J., and Gómez-Benito, M. (2013). Numerical modelling of the angiogenesis process in wound contraction. Biomechanics and modeling in mechanobiology 12, 349–360.

24. Valero, C., Javierre, E., García-Aznar, J., Gómez-Benito, M., and Menzel, A. (2015). Modeling of anisotropic wound healing. Journal of the Mechanics and Physics of Solids 79, 80–91.

25. Flegg, J.A., Menon, S.N., Maini, P.K., and McElwain, D.S. (2015). On the mathematical modeling of wound healing angiogenesis in skin as a reaction-transport process. Frontiers in physiology 6, 262.

26. Garbey, M., Salmon, R., Thanoon, D., and Bass, B.L. (2013). Multiscale modeling and distributed computing to predict cosmesis outcome after a lumpectomy. Journal of Computational Physics 244, 321–335.

27. Salmon, R., Garbey, M., Moore, L.W., and Bass, B.L. (2015). Interrogating a multifactorial model of breast conserving therapy with clinical data. Plos One 10, e0125006.

28. Vavourakis, V., Eiben, B., Hipwell, J.H., Williams, N.R., Keshtgar, M., and Hawkes, D.J. (2016). Multiscale mechano-biological finite element modelling of oncoplastic breast surgery—numerical study towards surgical planning and cosmetic outcome prediction. PloS One 11, e0159766.

29. Harbin, Z., Sohutskay, D., Vanderlaan, E., Fontaine, M., Mendenhall, C., Fisher, C., Voytik-Harbin, S., and Tepole, A.B. (2023). Computational mechanobiology model evaluating healing of postoperative cavities following breast-conserving surgery. Computers in Biology and Medicine 165, 107342.

30. Harbin, Z., Fisher, C., Voytik-Harbin, S., and Tepole, A.B. (2026). Computational modeling of patient-specific healing and deformation outcomes following breast-conserving surgery based on mri data. Annals of Biomedical Engineering 54, 495–513.

31. Fisher, B., Anderson, S., Bryant, J., Margolese, R.G., Deutsch, M., Fisher, E.R., Jeong, J.H., and Wolmark, N. (2002). Twenty-year follow-up of a randomized trial comparing total mastectomy, lumpectomy, and lumpectomy plus irradiation for the treatment of invasive breast cancer. New England Journal of Medicine 347, 1233–1241.

32. Olivotto, I.A., Lesperance, M.L., Truong, P.T., Nichol, A., Berrang, T., Tyldesley, S., Germain, F., Speers, C., Wai, E., Holloway, C. et al. (2009). Intervals longer than 20 weeks from breast-conserving surgery to radiation therapy are associated with inferior outcome for women with early-stage breast cancer who are not receiving chemotherapy. Journal of clinical oncology 27, 16–23.

33. Dahlbäck, C., Manjer, J., Rehn, M., and Ringberg, A. (2016). Determinants for patient satisfaction regarding aesthetic outcome and skin sensitivity after breast-conserving surgery. World Journal of Surgical Oncology 14, 1–11.

34. Brands-Appeldoorn, A., Thomma, R., Janssen, L., Maaskant-Braat, A., Tjan-Heijnen, V., and Roumen, R. (2022). Factors related to patient-reported cosmetic outcome after breast-conserving therapy for breast cancer. Breast Cancer Research and Treatment 191, 545–552.

35. Beadle, G.F., Silver, B., Botnick, L., Hellman, S., and Harris, J.R. (1984). Cosmetic results following primary radiation therapy for early breast cancer. Cancer 54, 2911–2918.

36. Roy, S., Biswas, S., Khanna, S., Gordillo, G., Bergdall, V., Green, J., Marsh, C.B., Gould, L.J., and Sen, C.K. (2009). Characterization of a preclinical model of chronic ischemic wound. Physiological genomics 37, 211–224.

37. Moor, A.N., Tummel, E., Prather, J.L., Jung, M., Lopez, J.J., Connors, S., and Gould, L.J. (2014). Consequences of age on ischemic wound healing in rats: altered antioxidant activity and delayed wound closure. Age 36, 733–748.

38. Harbin, Z.J., Fisher, C.S., Morrison, R.A., Gomez, H., Voytik-Harbin, S., and Buganza Tepole, A.B. (2026). Capillary network generation framework for estimating volumetric capillary density from histological vascular measurements. bioRxiv. URL: https://www.biorxiv.org/content/early/2026/07/13/2026.07.10.737824. doi: 10.64898/2026.07.10.737824. arXiv:https://www.biorxiv.org/content/early/2026/07/13/2026.07.10.737824.full.pdf

39. Nelson, T.R., Cerviño, L.I., Boone, J.M., and Lindfors, K.K. (2008). Classification of breast computed tomography data. Medical Physics 35, 1078–1086.

40. Valeta-Magara, A., Hatami, R., Axelrod, D., Roses, D.F., Guth, A., Formenti, S.C., and Schneider, R.J. (2015). Pro-oncogenic cytokines and growth factors are differentially expressed in the post-surgical wound fluid from malignant compared to benign breast lesions. SpringerPlus 4, 1–11.

41. Jeffrey, S.S., Goodson, W.H., Ikeda, D.M., Birdwell, R.L., and Bogetz, M.S. (1995). Axillary lymphadenectomy for breast cancer without axillary drainage. Archives of Surgery 130, 909–913.

42. Nissen, N.N., Polverini, P., Koch, A.E., Volin, M.V., Gamelli, R.L., and DiPietro, L.A. (1998). Vascular endothelial growth factor mediates angiogenic activity during the proliferative phase of wound healing. The American journal of pathology 152, 1445.

43. Gupta, S., Mujawdiya, P., Maheshwari, G., and Sagar, S. (2022). Dynamic role of oxygen in wound healing: a microbial, immunological, and biochemical perspective. Archives of Razi Institute 77, 513.

44. Remensnyder, J., and Majno, G. (1968). Oxygen gradients in healing wounds. The American Journal of Pathology 52, 301.

45. Tang, D., Yan, T., Zhang, J., Jiang, X., Zhang, D., and Huang, Y. (2017). Notch1 signaling contributes to hypoxia-induced high expression of integrin β1 in keratinocyte migration. Scientific reports 7, 43926.

46. Mirza, R., DiPietro, L.A., and Koh, T.J. (2009). Selective and specific macrophage ablation is detrimental to wound healing in mice. The American journal of pathology 175, 2454–2462.

47. Michalczyk, E.R., Chen, L., Fine, D., Zhao, Y., Mascarinas, E., Grippo, P.J., and DiPietro, L.A. (2018). Pigment epithelium-derived factor (pedf) as a regulator of wound angiogenesis. Scientific Reports 8, 11142.

48. Witte, M.B., and Barbul, A. (1997). General principles of wound healing. Surgical Clinics of North America 77, 509–528.

49. Okonkwo, U.A., Chen, L., Ma, D., Haywood, V.A., Barakat, M., Urao, N., and DiPietro, L.A. (2020). Compromised angiogenesis and vascular integrity in impaired diabetic wound healing. PloS one 15, e0231962.

50. Bonilla, E.V., Chai, K., and Williams, C. (2007). Multi-task gaussian process prediction. Advances in neural information processing systems 20.

51. Vos, E., Koppert, L., van Lankeren, W., Verhoef, C., Koerkamp, B.G., and Hunink, M. (2018). A preliminary prediction model for potentially guiding patient choices between breast conserving surgery and mastectomy in early breast cancer patients; a dutch experience. Quality of Life Research 27, 545–553.

52. Gardfjell, A., Dahlbäck, C., and Åhsberg, K. (2019). Patient satisfaction after unilateral oncoplastic volume displacement surgery for breast cancer, evaluated with the breast-q™. World Journal of Surgical Oncology 17, 1–13.

53. Veiga, D.F., Veiga-Filho, J., Ribeiro, L.M., Archangelo-Junior, I., Balbino, P.F., Caetano, L.V., Novo, N.F., and Ferreira, L.M. (2010). Quality-of-life and self-esteem outcomes after oncoplastic breast-conserving surgery [outcomes article]. Plastic and Reconstructive Surgery 125, 811–817.

54. Haubner, F., Ohmann, E., Pohl, F., Strutz, J., and Gassner, H.G. (2012). Wound healing after radiation therapy: review of the literature. Radiation oncology 7, 162.

55. Nunez-Alvarez, L., Ledwon, J.K., Applebaum, S., Progri, B., Han, T., Laudo, J., Tac, V., Gosain, A.K., and Tepole, A.B. (2024). Tissue expansion mitigates radiation-induced skin fibrosis in a porcine model. Acta Biomaterialia 189, 427–438.

56. Lawrenson, R., Lao, C., Stanley, J., Campbell, I., Krebs, J., Meredith, I., Koea, J., Teng, A., Sika-Paotonu, D., Stairmand, J. et al. (2023). Impact of diabetes on surgery and radiotherapy for breast cancer. Breast cancer research and treatment 199, 305–314.

57. Sherratt, J.A., and Murray, J. (1991). Mathematical analysis of a basic model for epidermal wound healing. Journal of Mathematical Biology 29, 389–404.

58. Urao, N., Okonkwo, U.A., Fang, M.M., Zhuang, Z.W., Koh, T.J., and DiPietro, L.A. (2016). Microct angiography detects vascular formation and regression in skin wound healing. Microvascular research 106, 57–66.

59. Bilionis, I., and Zabaras, N. (2014). Solution of inverse problems with limited forward solver evaluations: a bayesian perspective. Inverse Problems 30, 015004.

60. Rasmussen, C.E., and Williams, C.K. (2004). Gaussian processes in machine learning. Lecture Notes in Computer Science 3176, 63–71.

61. Dürichen, R., Pimentel, M.A., Clifton, L., Schweikard, A., and Clifton, D.A. (2014). Multi-task gaussian process models for biomedical applications. In IEEE-EMBS international Conference on Biomedical and health informatics (BHI). IEEE pp. 492–495.

62. Dürichen, R., Pimentel, M.A., Clifton, L., Schweikard, A., and Clifton, D.A. (2014). Multitask gaussian processes for multivariate physiological time-series analysis. IEEE transactions on biomedical engineering 62, 314–322.

63. Feng, X., Tonnesen, M.G., Mousa, S.A., and Clark, R.A. (2013). Fibrin and collagen differentially but synergistically regulate sprout angiogenesis of human dermal microvascular endothelial cells in 3-dimensional matrix. International journal of cell biology 2013, 231279.

64. Schindelin, J., Rueden, C.T., Hiner, M.C., and Eliceiri, K.W. (2015). The imagej ecosystem: An open platform for biomedical image analysis. Molecular reproduction and development 82, 518–529.

65. Gardner, J., Pleiss, G., Weinberger, K.Q., Bindel, D., and Wilson, A.G. (2018). Gpytorch: Blackbox matrix-matrix gaussian process inference with gpu acceleration. Advances in neural information processing systems 31.

66. Foreman-Mackey, D., Hogg, D.W., Lang, D., and Goodman, J. (2013). emcee: The MCMC Hammer. 125, 306. doi: 10.1086/670067. arXiv:1202.3665.

67. Buganza Tepole, A., and Kuhl, E. (2016). Computational modeling of chemo-bio-mechanical coupling: a systems-biology approach toward wound healing. Computer Methods in Biomechanics and Biomedical Engineering 19, 13–30.

68. Tepole, A.B. (2017). Computational systems mechanobiology of wound healing. Computer Methods in Applied Mechanics and Engineering 314, 46–70.

69. Sohutskay, D.O., Tepole, A.B., and Voytik-Harbin, S.L. (2021). Mechanobiological wound model for improved design and evaluation of collagen dermal replacement scaffolds. Acta Biomaterialia 135, 368–382.

70. TK, H. (1972). Role of oxygen in repair processes. Acta Chir Scand 138, 109–110.

71. Sohutskay, D.O., Puls, T.J., and Voytik-Harbin, S.L. (2020). Collagen self-assembly: biophysics and biosignaling for advanced tissue generation. Multi-scale Extracellular Matrix Mechanics and Mechanobiology pp. 203–245.

72. Cohen, I.K., Die-gelmann, R.F., Lindblad, W.J., and Hugo, N.E. (1992). Wound healing: biochemical and clinical aspects. Plastic and Reconstructive Surgery 90, 926.

73. Guyton, A.C., and Hall, J.E. (2021). Guyton and Hall Textbook of Medical Physiology. 14 ed.. Philadelphia, PA: Elsevier.

74. DiPietro, L.A. (2016). Angiogenesis and wound repair: when enough is enough. Journal of Leucocyte Biology 100, 979–984.

75. Distler, O., Distler, J.H., Scheid, A., Acker, T., Hirth, A., Rethage, J., Michel, B.A., Gay, R.E., Müller-Ladner, U., Matucci-Cerinic, M. et al. (2004). Uncontrolled expression of vascular endothelial growth factor and its receptors leads to insufficient skin angiogenesis in patients with systemic sclerosis. Circulation research 95, 109–116.

76. Steinbrech, D.S., Longaker, M.T., Mehrara, B.J., Saadeh, P.B., Chin, G.S., Gerrets, R.P., Chau, D.C., Rowe, N.M., and Gittes, G.K. (1999). Fibroblast response to hypoxia: the relationship between angiogenesis and matrix regulation. Journal of surgical Research 84, 127–133.

77. Steinbrech, D.S., Mehrara, B.J., Chau, D., Rowe, N.M., Chin, G., Lee, T., Saadeh, P.B., Gittes, G.K., and Longaker, M.T. (1999). Hypoxia upregulates vegf production in keloid fibroblasts. Annals of plastic surgery 42, 514–520.

78. Wu, W.K., Llewellyn, O.P., Bates, D.O., Nicholson, L.B., and Dick, A.D. (2010). Il-10 regulation of macrophage vegf production is dependent on macrophage polarisation and hypoxia. Immunobiology 215, 796–803.

79. Dannhauser, D., Rossi, D., De Gregorio, V., Netti, P.A., Terrazzano, G., and Causa, F. (2022). Single cell classification of macrophage subtypes by label-free cell signatures and machine learning. Royal Society Open Science 9.

80. Vaupel, P., Mayer, A., Briest, S., and Höckel, M. (2005). Hypoxia in breast cancer: role of blood flow, oxygen diffusion distances, and anemia in the development of oxygen depletion. Oxygen Transport to Tissue XXVI pp. 333–342.

## References

1. Puls, T.J., Fisher, C.S., Cox, A., Plantenga, J.M., McBride, E.L., Anderson, J.L., Goergen, C.J., Bible, M., Moller, T., and Voytik-Harbin, S.L. (2021). Regenerative tissue filler for breast conserving surgery and other soft tissue restoration and reconstruction needs. Scientific Reports 11, 2711.

2. Sohutskay, D.O., Puls, T.J., and Voytik-Harbin, S.L. (2020). Collagen self-assembly: biophysics and biosignaling for advanced tissue generation. Multi-scale Extracellular Matrix Mechanics and Mechanobiology pp. 203–245.

3. Harbin, Z., Sohutskay, D., Vanderlaan, E., Fontaine, M., Mendenhall, C., Fisher, C., Voytik-Harbin, S., and Tepole, A.B. (2023). Computational mechanobiology model evaluating healing of postoperative cavities following breast-conserving surgery. Computers in Biology and Medicine 165, 107342.

4. Olsen, L., Sherratt, J.A., and Maini, P.K. (1995). A mechanochemical model for adult dermal wound contraction and the permanence of the contracted tissue displacement profile. Journal of Theoretical Biology 177, 113–128.

5. TK, H. (1972). Role of oxygen in repair processes. Acta Chir Scand 138, 109–110.

6. Tepole, A.B. (2017). Computational systems mechanobiology of wound healing. Computer Methods in Applied Mechanics and Engineering 314, 46–70.

7. Valero, C., Javierre, E., García-Aznar, J.M., and Gómez-Benito, M.J. (2014). A cell-regulatory mechanism involving feedback between contraction and tissue formation guides wound healing progression. PloS One 9, e92774.

8. Cumming, B.D., McElwain, D., and Upton, Z. (2010). A mathematical model of wound healing and subsequent scarring. Journal of The Royal Society Interface 7, 19–34.

9. Koppenol, D.C., Vermolen, F.J., Niessen, F.B., van Zuijlen, P.P., and Vuik, K. (2017). A mathematical model for the simulation of the formation and the subsequent regression of hypertrophic scar tissue after dermal wounding. Biomechanics and Modeling in Mechanobiology 16, 15–32.

10. Murphy, K.E., Hall, C.L., Maini, P.K., McCue, S.W., and McElwain, D.S. (2012). A fibrocontractive mechanochemical model of dermal wound closure incorporating realistic growth factor kinetics. Bulletin of Mathematical Biology 74, 1143–1170.

11. Cohen, I.K., Die-gelmann, R.F., Lindblad, W.J., and Hugo, N.E. (1992). Wound healing: biochemical and clinical aspects. Plastic and Reconstructive Surgery 90, 926.

12. Tepole, A.B., Ploch, C.J., Wong, J., Gosain, A.K., and Kuhl, E. (2011). Growing skin: a computational model for skin expansion in reconstructive surgery. Journal of the Mechanics and Physics of Solids 59, 2177–2190.

13. Laurent, G. (1987). Dynamic state of collagen: pathways of collagen degradation in vivo and their possible role in regulation of collagen mass. American Journal of Physiology-Cell Physiology 252, C1–C9.

14. Vaupel, P., Mayer, A., Briest, S., and Höckel, M. (2005). Hypoxia in breast cancer: role of blood flow, oxygen diffusion distances, and anemia in the development of oxygen depletion. Oxygen Transport to Tissue XXVI pp. 333–342.

15. Xue, C., Friedman, A., and Sen, C.K. (2009). A mathematical model of ischemic cutaneous wounds. Proceedings of the National Academy of Sciences 106, 16782–16787.

16. Guyton, A.C., and Hall, J.E. (2021). Guyton and Hall Textbook of Medical Physiology. 14 ed.. Philadelphia, PA: Elsevier.

17. Feng, X., Tonnesen, M.G., Mousa, S.A., and Clark, R.A. (2013). Fibrin and collagen differentially but synergistically regulate sprout angiogenesis of human dermal microvascular endothelial cells in 3-dimensional matrix. International journal of cell biology 2013, 231279.

18. Streeter, I., and Cheema, U. (2011). Oxygen consumption rate of cells in 3d culture: the use of experiment and simulation to measure kinetic parameters and optimise culture conditions. Analyst 136, 4013–4019.

19. Chin, M.P., Schauer, D.B., and Deen, W.M. (2010). Nitric oxide, oxygen, and superoxide formation and consumption in macrophages and colonic epithelial cells. Chemical research in toxicology 23, 778–787.

20. Baker, E.A., Kumar, S., Melling, A.C., Whetter, D., and Leaper, D.J. (2008). Temporal and quantitative profiles of growth factors and metalloproteinases in acute wound fluid after mastectomy. Wound Repair and Regeneration 16, 95–101.

21. Steinbrech, D.S., Longaker, M.T., Mehrara, B.J., Saadeh, P.B., Chin, G.S., Gerrets, R.P., Chau, D.C., Rowe, N.M., and Gittes, G.K. (1999). Fibroblast response to hypoxia: the relationship between angiogenesis and matrix regulation. Journal of surgical Research 84, 127–133.

22. Steinbrech, D.S., Mehrara, B.J., Chau, D., Rowe, N.M., Chin, G., Lee, T., Saadeh, P.B., Gittes, G.K., and Longaker, M.T. (1999). Hypoxia upregulates vegf production in keloid fibroblasts. Annals of plastic surgery 42, 514–520.

23. Distler, O., Distler, J.H., Scheid, A., Acker, T., Hirth, A., Rethage, J., Michel, B.A., Gay, R.E., Müller-Ladner, U., Matucci-Cerinic, M. et al. (2004). Uncontrolled expression of vascular endothelial growth factor and its receptors leads to insufficient skin angiogenesis in patients with systemic sclerosis. Circulation research 95, 109–116.

24. Wu, W.K., Llewellyn, O.P., Bates, D.O., Nicholson, L.B., and Dick, A.D. (2010). Il-10 regulation of macrophage vegf production is dependent on macrophage polarisation and hypoxia. Immunobiology 215, 796–803.

25. Sanchez, B., Li, L., Dulong, J., Aimond, G., Lamartine, J., Liu, G., and Sigaudo-Roussel, D. (2019). Impact of human dermal microvascular endothelial cells on primary dermal fibroblasts in response to inflammatory stress. Frontiers in Cell and Developmental Biology 7, 44.

26. Bianchi, A., Painter, K.J., and Sherratt, J.A. (2016). Spatio-temporal models of lymphangiogenesis in wound healing. Bulletin of mathematical biology 78, 1904–1941.

27. Stockmann, C., Kirmse, S., Helfrich, I., Weidemann, A., Takeda, N., Doedens, A., and Johnson, R.S. (2011). A wound size–dependent effect of myeloid cell–derived vascular endothelial growth factor on wound healing. Journal of Investigative Dermatology 131, 797–801.

